# Generation and network analysis of an RNA-seq transcriptional atlas for the rat

**DOI:** 10.1101/2021.11.07.467633

**Authors:** Kim M. Summers, Stephen J. Bush, Chunlei Wu, David A. Hume

## Abstract

The laboratory rat is an important model for biomedical research. To generate a comprehensive rat transcriptomic atlas, we curated and down-loaded 7700 rat RNA-seq datasets from public repositories, down-sampled them to a common depth and quantified expression. Data from 590 rat tissues and cells, averaged from each Bioproject, can be visualised and queried at http://biogps.org/ratatlas. Gene correlation network (GCN) analysis revealed clusters of transcripts that were tissue or cell-type restricted and contained transcription factors implicated in lineage determination. Other clusters were enriched for transcripts associated with biological processes. Many of these clusters overlap with previous data from analysis of other species whilst some (e.g. expressed specifically in immune cells, retina/pineal gland, pituitary and germ cells) are unique to these data. GCN on large subsets of the data related specifically to liver, nervous system, kidney, musculoskeletal system and cardiovascular system enabled deconvolution of cell-type specific signatures. The approach is extensible and the dataset can be used as a point of reference from which to analyse the transcriptomes of cell types and tissues that have not yet been sampled. Sets of strictly co-expressed transcripts provide a resource for critical interpretation of single cell RNA-seq data.

## INTRODUCTION

In the year of the rat (2020), the Rat Genome Database (RGD) celebrated 20 years of development (1). Those 20 years saw completion of the draft genome (2). Around 90% of protein-coding genes had an inferred 1:1 ortholog in humans. Subsequent technology advances allowing the sequencing of multiple inbred strains including several with disease-associated alleles (3). Szpirer (4) catalogued more than 350 rat genes where rat lines with natural or introduced variants provide models for human disease.

Analysis of transcriptional regulation in human and mouse has been driven by large consortium projects such as GTEX (5) and FANTOM (6) and there are many on-line resources for these species. Multi-tissue transcriptional atlas projects have also been published for other species including chicken, sheep, buffalo, pig and goat (7-11). Although it was once suggested that guilt-by-association is the exception rather than the rule in gene regulatory networks (12), the principle is now very well-established. Genes associated with specific organs, cell types, organelles and pathways (e.g. the cell cycle, protein synthesis, oxidative phosphorylation/mitochondria) are stringently co-expressed along with the transcription factors that regulate them (5,6,8,13-18). An extension of the principle of co-regulated expression is that it is possible to extract signatures of specific cell types, for example the stromal component of tumors (19) or resident tissue macrophages (20) based upon analysis of a large number of samples in which their relative abundance is variable.

The functional annotation of the rat genome is still a work in progress. Many rat genes in Ensembl are described as “novel rat gene” and annotated solely by a gene number. Transcriptional regulation has evolved rapidly amongst mammalian species (21,22). Even where there is 1:1 orthology at the level of protein-coding sequence and conservation of synteny with other mammals the expression may not be conserved. Two substantial studies have contributed to annotation of the rat transcriptome through RNA-seq analysis of a partly-overlapping set of major rat organs (23,24). Long read RNA sequencing has also contributed to refinement of rat transcriptome annotation (25). Because of the extensive use of the rat as a model in biomedical research, there are thousands of RNA-seq datasets in the public domain from isolated cells and tissues in various states of activation that could provide an additional resource for functional annotation. By combining random library down-sizing to reduce sampling bias and the high-speed ‘pseudo-aligner’ Kallisto (26) to quantify expression, we previously established a pipeline [7, 11] to enable meta-analysis of published RNA-seq data. Here we have used this pipeline to produce an extended expression atlas for the laboratory rat. To demonstrate the robustness of the integrated data we have carried out network analysis to identify sets of co-expressed transcripts. The dataset is downloadable and the pipeline is extensible to allow inclusion of additional data and regeneration of the network as new RNA-seq data becomes available.

## METHODS

### Selecting samples for an expression atlas of the rat

To create a comprehensive expression atlas for the rat we first downloaded the daily-updated NCBI BioProject summary file from ftp://ftp.ncbi.nlm.nih.gov/bioproject/summary.txt (obtained 19th July 2021) and parsed it to obtain all BioProjects with taxonomy ID 10116 (*Rattus norvegicus*) and a data type of ‘transcriptome or gene expression’, supplementing this list by manually searching NCBI Geo and NCBI PubMed for the keywords “RNA-seq AND rat”. BioProjects were selected to extend the diversity of tissues, cells and states from two existing rat transcriptomic atlases that analyse gene expression in a subset of major rat tissues (23,24). For each BioProject, we automatically extracted the associated metadata using pysradb v1.0.1 (27) with parameter ‘--detailed’, or by manual review. Metadata for each BioProject, indicating (where available) the breed/strain, sex, age, tissue/cell type extracted, and experimental condition (for example, treatment or control) are detailed in **Table S1**, which includes both the data downloaded via the pipeline and additional information retrieved manually from the ENA record, NCBI BioProject record and cited publications. For incorporation into the expression atlas, we required that all samples have, at minimum, tissue/cell type recorded. Overall, the input to the atlas comprised 7682 samples from 363 BioProjects.

### Quantifying gene expression for the atlas

For each library, expression was quantified using Kallisto v0.44.0 (26) as described in detail in previous studies on other species (7-9,20). Kallisto quantifies expression at the transcript level, as transcripts per million (TPM), by building an index of k-mers from a set of reference transcripts and then ‘pseudo-aligning’ reads to it, matching k-mers in the reads to k-mers in the index. Transcript-level TPM estimates were then summed to give gene-level TPM. To create the reference transcriptomic index, we performed a non-redundant integration of the set of Ensembl v98 Rnor6.0 protein-coding cDNAs (http://ftp.ensembl.org/pub/release-98/fasta/rattus_norvegicus/cdna/Rattus_norvegicus.Rnor_6.0.cdna.all.fa.gz, accessed 24^th^ November 2019; n = 31,715 transcripts) and the set of 69,440 NCBI mRNA RefSeqs (https://ftp.ncbi.nlm.nih.gov/genomes/refseq/vertebrate_mammalian/Rattus_norvegicus/all_assembly_versions/suppressed/GCF_000001895.5_Rnor_6.0/GCF_000001895.5_Rnor_6.0_rna.fna.gz, accessed 24^th^ November 2019), as previously described (7). The purpose of the integration was to include transcripts that had not already been assigned Ensembl transcript IDs and whose sequence was not already present in the Ensembl release (under any identifier). RefSeq mRNAs incorporate untranslated regions (UTRs) and so could encapsulate an Ensembl CDS. The trimmed UTRs from each mRNA were generated excluding all sequence outside the longest ORF. In total, the reference transcriptome comprised 71,074 transcripts, representing 25,013 genes. Using this reference, expression was quantified for 7682 publicly-archived paired-end Illumina RNA-seq libraries. The Bioprojects are summarised in **Table S1**. Prior to expression quantification, and for the purpose of minimising variation between samples, we randomly downsampled all libraries to 10 million reads, 5 times each, using seqtk v1.2 (https://github.com/lh3/seqtk, downloaded 4^th^ June 2018). Expression level was then taken to be the median TPM across the 5 downsampled replicates.

The final expression atlas details the median downsampled TPM per gene, averaged for tissue, age, and BioProject. As in previous projects for other species (7-11) the full dataset of 590 averaged expression data from cells and tissues is displayed on BioGPS (28,29) at biogps.org/ratatlas to enable comparative analysis across species. The full processed primary dataset and the averaged data is available for download at an Institutional Repository (https://doi.org/10.5287/bodleian:Am9akye72). The latter is a comma-separated text file, which can be directly loaded into the network analysis software used herein or alternatives such as Gephi (gephi.org) or Cytoscape (cytoscape.org). This file can be easily supplemented by addition of further RNA-seq data processed in the same way. All scripts for generating the atlas are available at www.github.com/sjbush/expr_atlas.

### Network analysis and functional clustering of atlas samples

To examine the expression of genes across this wide range of tissues and cell types, the expression data were analysed using the network analysis tool BioLayout (derived from Biolayout *Express*^3D^ (30)), downloaded from http://biolayout.org. The same files can be uploaded into the recently-developed open source package, Graphia (https://graphia.app), which supports alternative clustering approaches and dynamic modification of parameters. The initial analysis used the values averaged by age and BioProject for each tissue. Subsequent analyses used individual values for samples of liver, musculoskeletal system, cardiovascular system, kidney and central nervous system. For each analysis, a sample to sample correlation matrix was initially constructed at the Pearson correlation coefficient (*r*) threshold necessary to include all samples in the analysis (shown in Results and figure legends). Pearson correlations were then calculated between all pairs of genes to produce a gene-to-gene correlation matrix of all genes correlated at *r* ≥ 0.75.

Gene co-expression networks (GCNs) were generated from the matrices, where nodes represent either samples or genes and edges represent correlations between nodes above the selected correlation threshold. For the sample-to-sample analyses (essentially analogous to a principal components analysis, PCA) an initial screen at the *r* value which entered all samples was performed, followed by subsequent analyses with a higher *r* value which removed outliers and revealed more substructure in the networks. For each gene-to-gene analysis an *r* value threshold of 0.75 was used for all analyses (**Figure S1**).

For the gene-to-gene networks, further analysis was performed to identify groups of highly connected genes within the overall topology of the network, using the Markov clustering algorithm (MCL) (31). The MCL is an algebraic bootstrapping process in which the number of clusters is not specified. A parameter called inflation effectively controls granularity. The choice of inflation value is empirical and is based in some measure on the predicted complexity of the dataset (31). The chosen inflation value was 2.2 for all analyses and only genes expressed at ≥ 10 TPM in at least one sample were included. Gene ontology (GO) terms and Reactome pathways were derived from the Gene Ontology Resource (http://geneontology.org, release of 18 August 2021) using PANTHER overrepresentation test (PANTHER release of 24 February 2021). The reference list used was *Rattus norvegicus* (all genes in database), the GO Ontology database was the release of 2 July 2021 (DOI: 10.5281/zenodo.5080993) and the Reactome pathway analysis used Reactome version 65, released 17 November 2020. These resources area all available at the Gene Ontology Resource (http://geneontology.org).

## RESULTS

### Samples in the atlas

7682 RNA-seq libraries, each with a unique SRA sample accession from 363 BioProjects, were obtained by the pipeline as described in Methods and used to create a global atlas of gene expression. Metadata for the individual BioProjects are summarised in **Table S1**. For comparative tissue analysis and the core atlas, expression across libraries was averaged by tissue, age and BioProject. This reduced the dataset to 590 different averaged samples of rat tissues and cells summarised in **Table S2A**. For a separate analysis of liver, kidney, musculoskeletal, cardiovascular and central nervous systems to extract tissue-specific co-expression signatures, individual RNA-seq datasets from within each BioProject were used.

### Network analysis of the rat transcriptome

Initially we performed a sample-to-sample correlation to assess whether there were likely to be batch effects resulting in outlier samples that were unrelated to tissue type. To include all 590 samples, it was necessary to use *r* ≥ 0.21. An image of the resulting network graph is shown in **Figure 1**. Since BioProjects tended to focus on one strain, age, sex and tissue/treatment, some BioProject specific clustering was expected. However, illustrating the robustness of the sampling and down-sizing approach, related tissues analysed in different BioProjects generally clustered together (compare Figure 1A where nodes are coloured by organ system and Figure 1B where they are coloured by BioProject). At a more stringent correlation coefficient threshold of 0.7, only 15 samples of relatively low connectivity were removed but the association of nodes by organ system rather than BioProject is more obvious (**Figure 1C and D**). No clear outliers or BioProject-specific clusters (batch effects) were identified so all averaged samples were included in the subsequent gene-centred network analysis. The threshold correlation coefficient was chosen to maximise the number of nodes (genes included) while minimising the number of edges (correlations between them) (**Figure S1**). At the optimal correlation coefficient of *r* ≥ 0.75, the graph contained 14,848 nodes (genes) connected by 1,152,325 edges.

**Figure 1.**
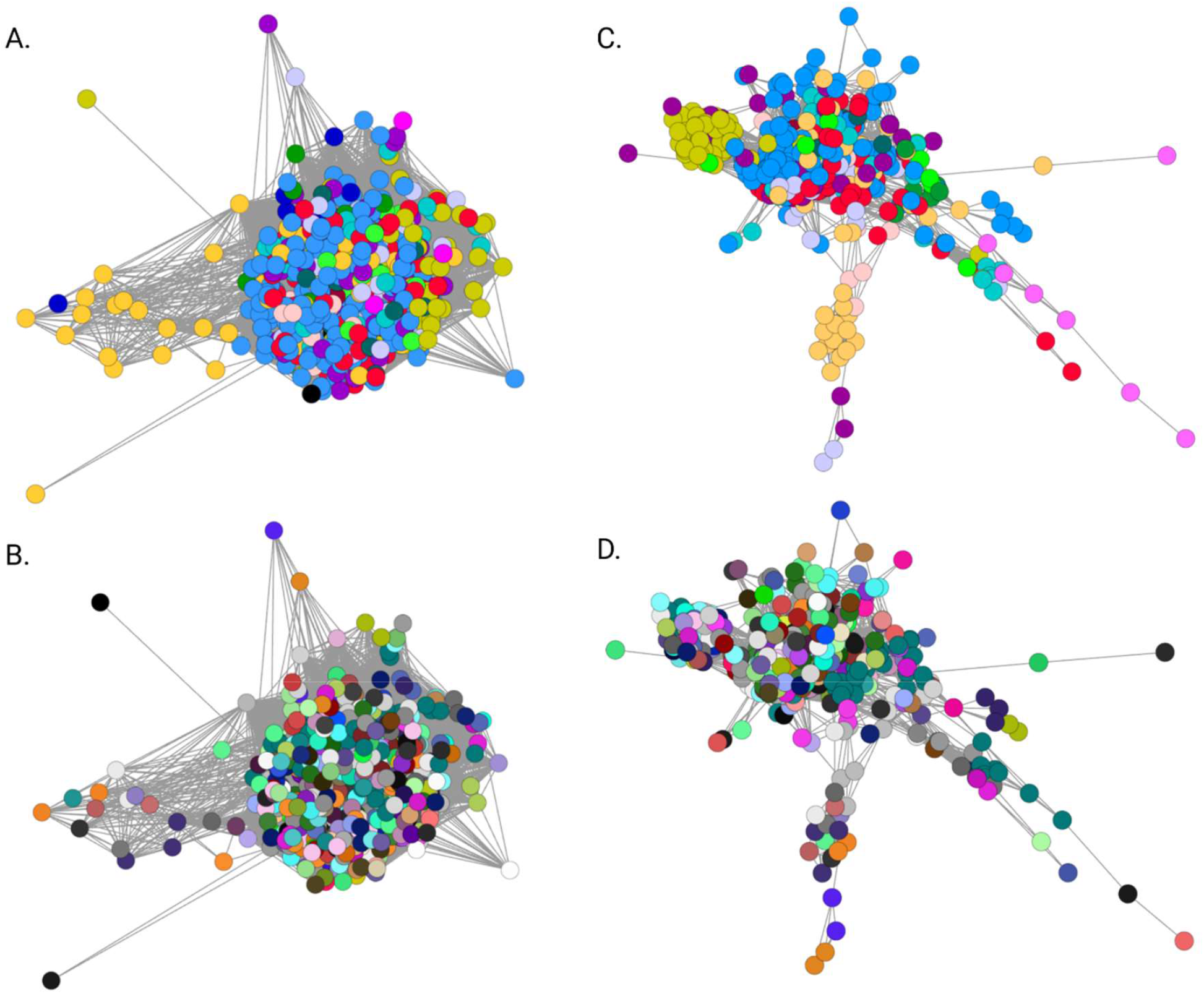
Sample to sample network graph for samples averaged by BioProject, age and tissue type. **A**. and **C**. Nodes coloured by organ system. Dark red – auditory system; light red, cardiovascular system, salmon, digestive system; orange, endocrine system; olive, liver; bright green, female reproductive system; teal, immune system; dark teal, integumentary system; dark green, male reproductive system; black, mixed tissues; light blue, nervous system; dark blue, primordia/early development; purple, renal system, pink, respiratory system; mauve, skeletomuscular system; grey, whole body (embryo). **B**. and **D**. Nodes coloured by BioProject. For **A**. and **B**. a correlation coefficient threshold of 0.21 was used; for **C**. and **D**, the threshold was 0.7.

**Table S2B** shows all of the clusters detected for transcripts with a minimum expression of ≥ 10TPM in at least one sample. By comparison to previous network analysis of mouse, human, pig, chicken, sheep and water buffalo transcriptomes (7-11) at this relatively stringent correlation coefficient, the much larger and more diverse rat transcriptomic dataset has a more fine-grained distribution with >1300 clusters having 2 nodes or more. In the published RNA-seq transcriptional atlas of 11 rat organs (32) which is included in the current data, around 40% of transcripts were expressed in all organs, in both sexes and at all development stages. In this larger set of averaged data, reflecting the much greater diversity of tissues and isolated cells sampled here, only 96 genes (0.38%) were detected above the 10 TPM minimal threshold in all 590 samples.

GO terms for clusters discussed below are included in **Table S2C**. Consistent with previous analysis, there are clusters that show no evidence of tissue-specificity but are clearly-enriched for genes involved in defined biological functions. For example, Cluster 11, Cluster 54 and Cluster 69 are associated with the cell cycle, DNA synthesis and repair. Cluster 41 is made up almost entirely of histone-encoding transcripts, likely due to incomplete removal of non-polyadenylated transcripts in some of the RNA-seq libraries. This cluster is not specific to any BioProject. The 18 transcripts within this cluster identified by LOCID also have provisional annotation as histones. Although this cluster is the product of a technical error, it also highlights the power of the clustering approach to extract signatures of co-expression.

**Table 1** summarises the expression patterns and biological processes associated with clusters of transcripts showing evidence of tissue or cell-type enrichment. The largest cluster of transcripts (Cluster 1), >1500 in total, is expressed almost exclusively in the testis. More than 500 of these transcripts are identified only by a LOCID, RGD or other uninformative annotation and many more are identified only by structural motif (for example 50 members of the Ccdc family, 35 undefined Fams, 20 testis-expressed (Tex) and 15 Tmem protein genes). The complexity of the testis transcriptome in all mammalian species has been widely recognised (reviewed in (33)). The set of testis-enriched transcripts with functional annotations encode proteins associated with meiosis, sperm differentiation, structure and motility and acrosomes. Unannotated genes are likely to involved in male fertility. For example, mutation of *Dlec1*, a putative tumor suppressor gene, was recently shown to cause male infertility in mice (34). LOC498675 is a predicted 1:1 ortholog of mouse testis-specific gene 1700102P08Rik, which is expressed in spermatocytes and is essential for male fertility (35,36). Smaller testis-enriched clusters include Cluster 29, which contains Sertoli cell markers such as *Aard* and *Tsx* (37,38), Cluster 72, which contains *Fshr* and the essential testis-specific transcription factor *Taf7l* ((39,40)) and Cluster 88, which includes the male-determining transcription factor *Sry*.

**Table 1.**
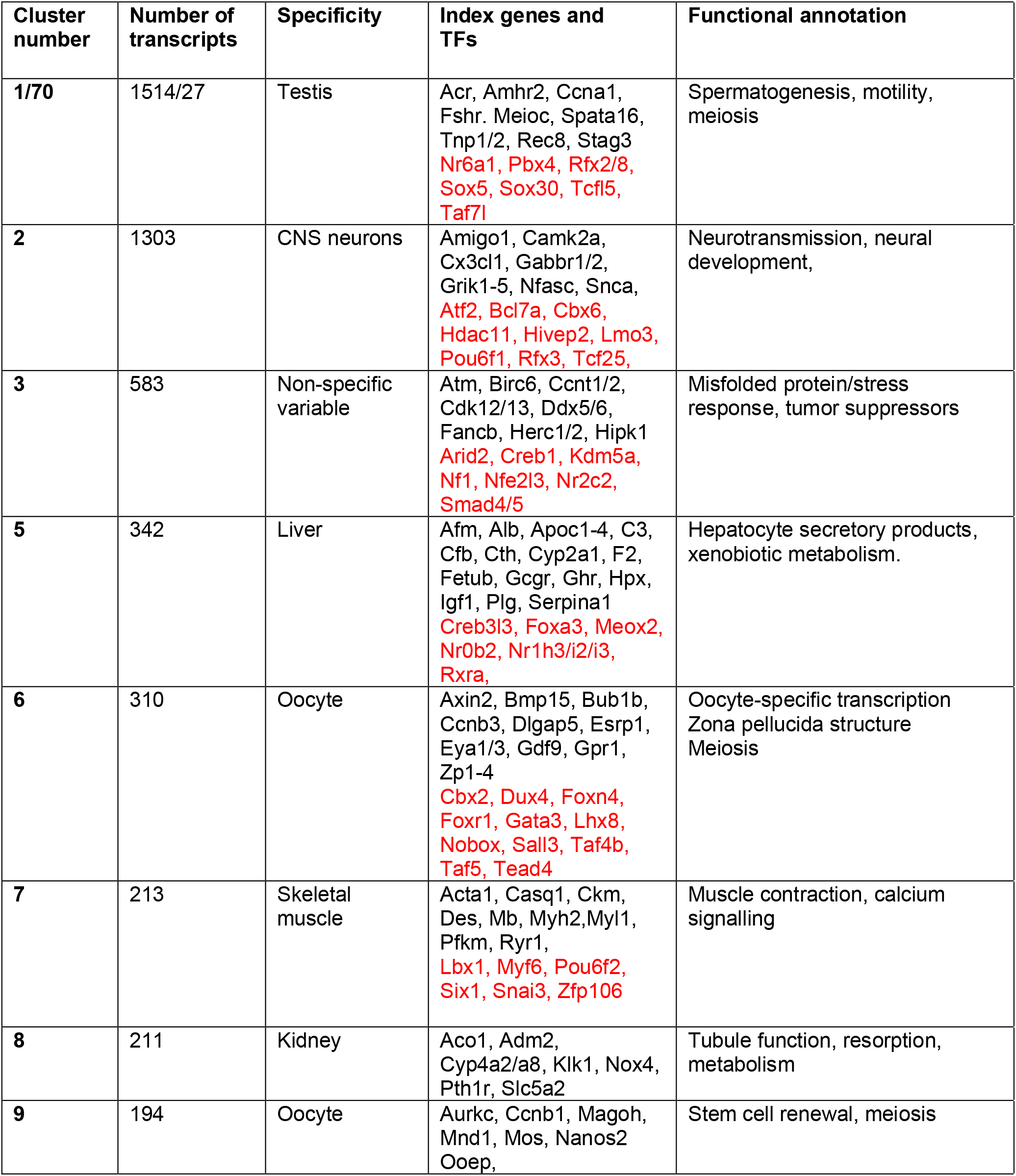
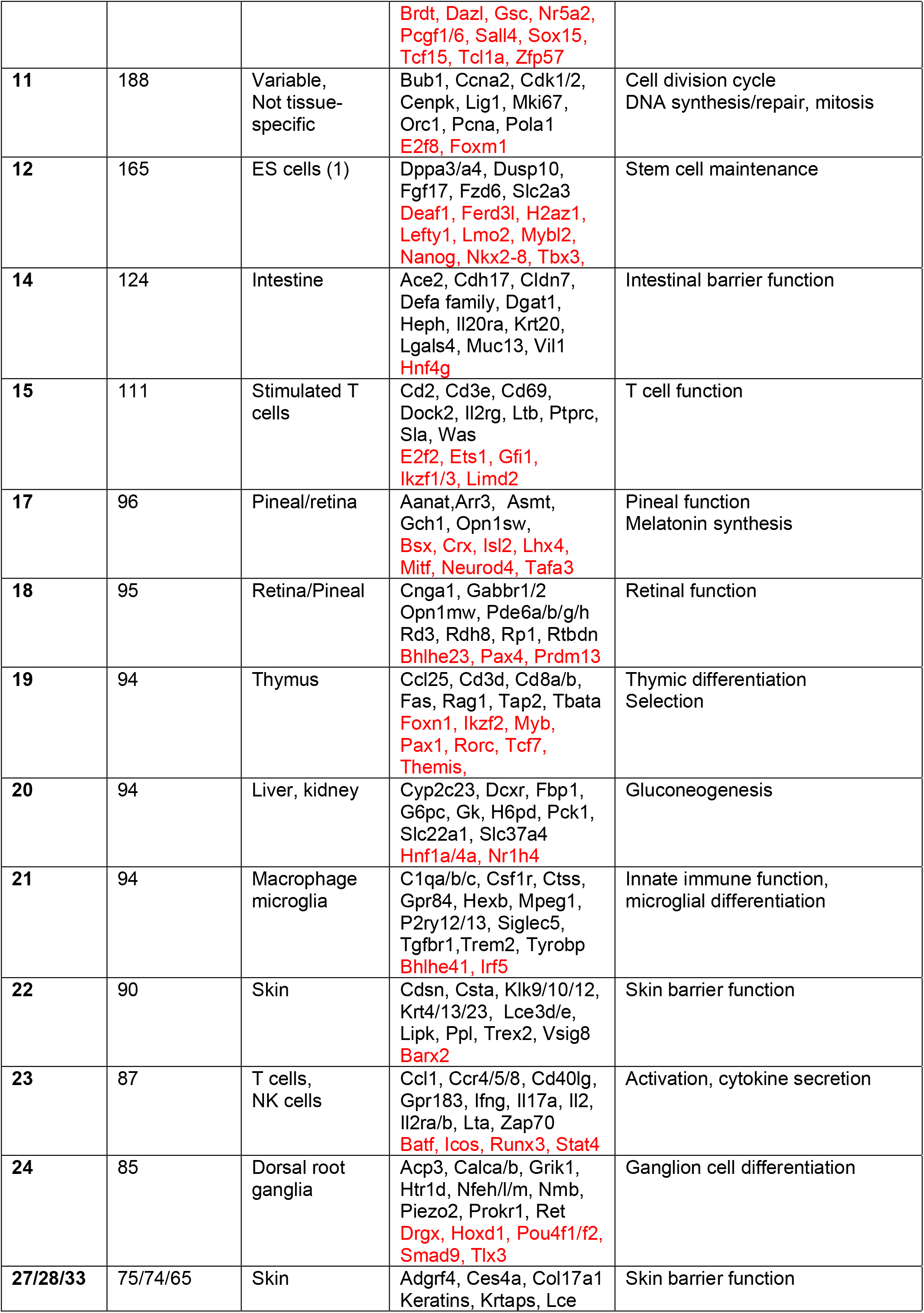
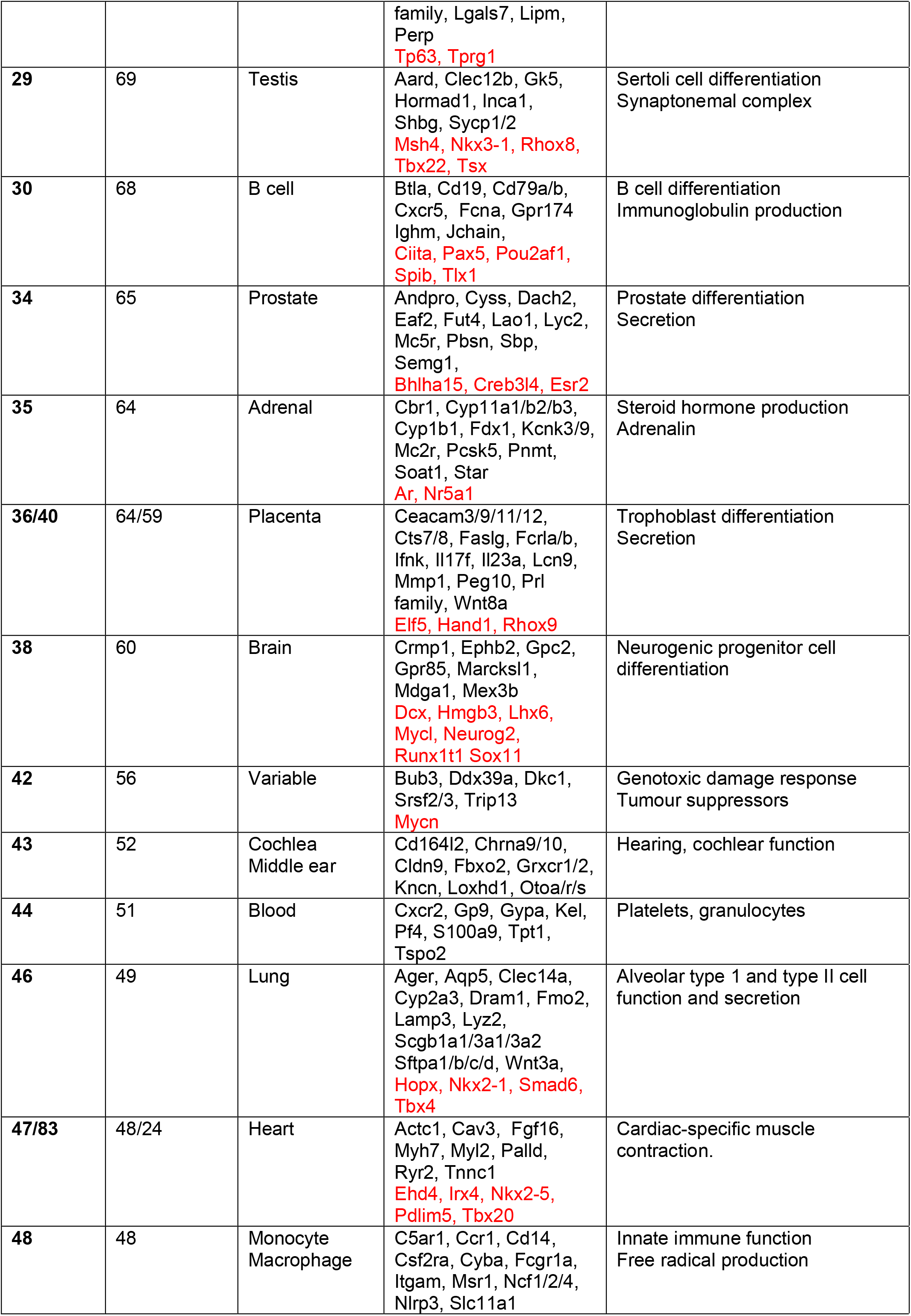
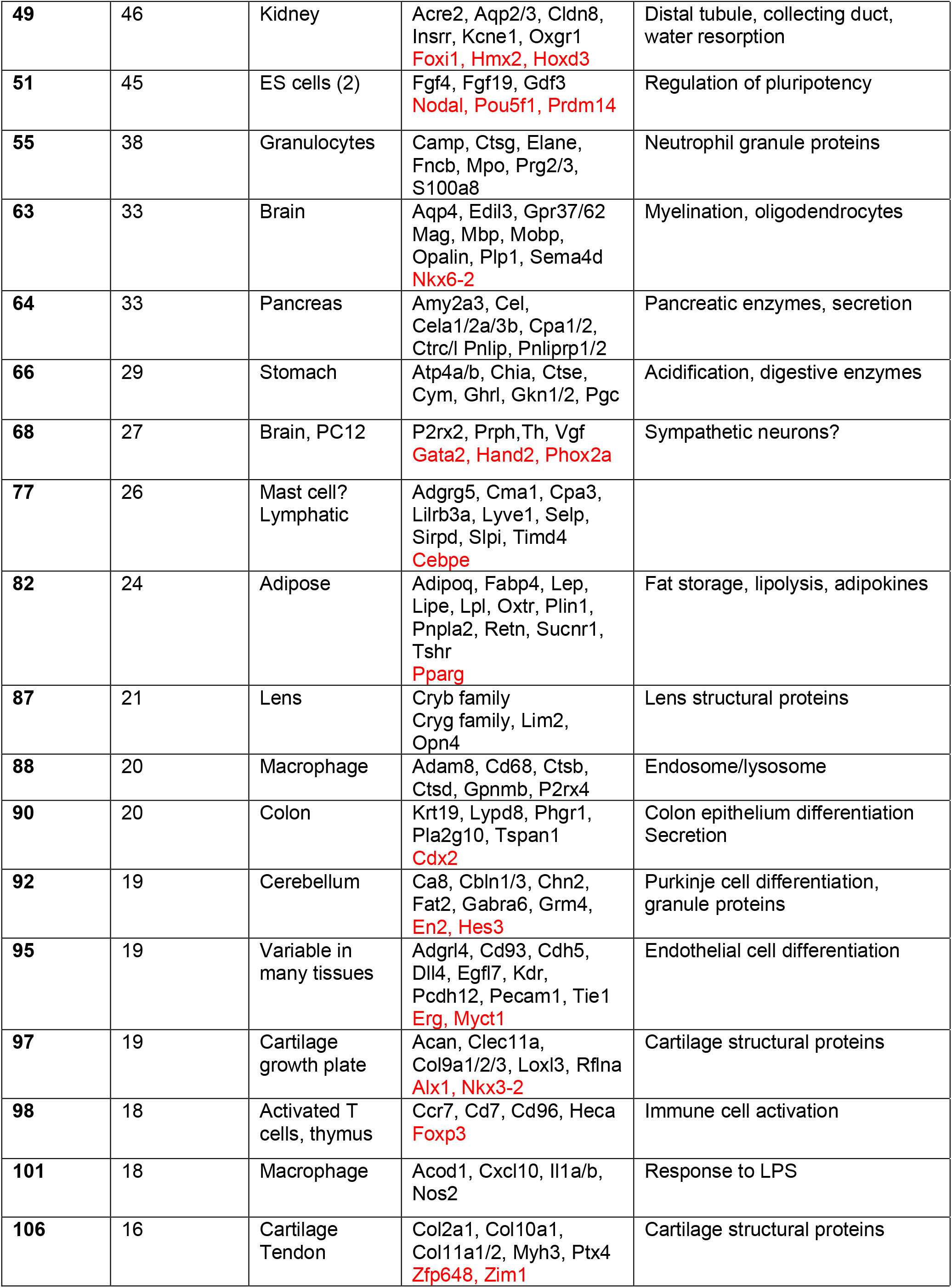
Gene expression clusters from rat tissues and cells. Clusters were generated at *r* ≥ 0.75 and MCL inflation value 2.2. Clusters of ≥ 40 nodes are shown. Selected transcripts encoding transcription factors are highlighted in red.

Clusters 17 and 18 contain transcripts expressed in both the retina and the pineal gland, both intimately involved in chronobiology and light sensing. Chang *et al*. (41) recently produced an aggregated resource describing the shared and divergent transcriptomes of these structures. Cluster 17 contains *Opn1sw*, the pineal-enriched transcription factor *Crx* and its target *Aanat* encoding the rate-limiting enzyme in melatonin synthesis (42). One unexpected inclusion in Cluster 17, enriched in pineal, is the transcript encoding the transcription factor MITF. *Mitf* in humans may be driven by as many as 7 distinct promoters including one used specifically by melanocytes. A unique transcription start site is shared by retinal pigment epithelial cells and pineal gland. *Mitf* over-expression in mouse pineal gland relative to other tissues has been noted previously (42,43) and in humans also *MITF* is most highly-expressed in pineal (http://biogps.org). However, whereas targets of MITF have been identified in melanocytes and many other cell types (44) and mutations impact many complex phenotypes in mice and humans, there appears to be no literature on its role in the pineal. To illustrate the utility of the data, in **Table S2D** we have reviewed the annotation of transcripts in Clusters 17 and 18 identified as LOCID. Several novel transcripts of unknown function (e.g. *Katnip* (*LOC361646*,aka KIAA0586; Talpid3), encoding a highly-conserved ciliary protein associated with the human genetic disease, Joubert syndrome (45) and *Lrtm1* (*LOC102547963*), a novel membrane protein) are also almost uniquely expressed in the human pineal gland (http://biogps.org)

Many small clusters are enriched in tissues, cell-types or activation states that were not analysed in the existing rat atlases or indeed in any previous atlas project in other species. They can be annotated based upon known markers. For example, Cluster 145 with 12 nodes contains transcripts encoding major secreted products of the pituitary (*Cga, Gh1, Fshb, Lhb, Tshb*) and the transcription factors that regulate their expression (*Pitx1, Six6, Tbx19*). Cluster 180 contains a subset of known immediate early genes (*Egr1,Fos,Jun*) mostly associated with isolated primary cells, and likely reflects cell activation during isolation or tissue processing (20). Other known genes in the immediate early class cluster separately, or not at all, because they are constitutively expressed by specific cell types. Similarly, groups of inducible genes in innate immune cells are all expressed by LPS-stimulated macrophages but divide into at least three clusters (Cluster 101, including *Il1a*; Cluster 112 including *Ifit2* and other interferon targets; Cluster 126 (including *Tnf*) because of expression by non-immune cells.

Other smaller clusters group genes that share functions. The large protocadherin family of cell adhesion molecules is broadly-divided into the clustered (α, β γ) and non-clustered (δ) subgroups (46). The δ protocadherins are predominantly expressed in the nervous system and indeed *Pcdh1, 8, 9, 20* are brain-restricted and part of the second largest cluster (**Cluster 2**). However, Cluster 81 includes *Pcdhb22* and16 members of the *Pcdhg* (A and B) families which are collectively enriched in the CNS but also widely expressed in other tissues. In addition, LOC108353166 within this cluster is annotated as protocadherin gamma-B2-like. Further members are more brain-restricted and grouped together in Cluster 250. 9/13 mitochondrially-encoded peptides group together in Cluster 212 whereas Clusters 61 and 76 group nuclear-encoded mitochondrial genes involved in the TCA cycle and oxidative phosphorylation (as expected, most highly-expressed in heart and kidney). Cluster 102 groups 18 transcripts encoding proteins involved in mitochondrial β oxidation of fatty acids. Several of the genes in this cluster are mutated in multiple acyl-CoA dehydrogenase deficiency (MADD, also known as glutaric aciduria type II) and related metabolic disorders (47). One additional gene involved in this pathway, *Etfb*, does not form part of a cluster. It is correlated with *Etfa* at *r* = 0.599 and with *Etfdh* at *r* = 0.527 but expressed at lower levels in certain tissues including the pineal gland.

Cluster 127, with 14 nodes, contains two markers of neurogenic cells (*Sstr2, Mpped1*; (48,49)) and a candidate regulator, *Tiam2* (50) and is otherwise made up of 11 brain-specific transcriptional regulators, each of which has been shown to be essential for neurogenesis and likely interacts with the others. Clusters 125 and 332 contains 20 genes encoding proteins that have all been implicated as molecular chaperones including multiple components of the TRIC chaperone complex (*Tcp1, Cct2,3,4,5*). Cluster 557 with only 4 nodes contains the oligodendrocyte transcription factors, *Olig1* and *Olig2*, as well as *Sox 8*, which has non-redundant function in oligodendrocyte differentiation (51). The fourth node in this cluster, LOC103692025, is predicted by the RGD to be an ortholog of *Lhfpl3* which in mouse is a marker of oligodendrocyte lineage commitment (52). The two calmodulin-encoding genes (*Calm1* and *Calm2*) are co-expressed (Cluster 673) as are three genes involved in cholesterol synthesis (*Fdft1, Hmgcr, Hmgcs1*) (Cluster 742). *Ins1* and *Ins2*, encoding insulin, are co-expressed with pancreatic polypeptide (*Ppy*) (Cluster 751) but not with glucagon (*Gcg*). Although *Ppy* is normally expressed by rare gamma cells in pancreatic islets, a recent study indicated that gamma cells can produce insulin following beta cell injury (53).

Each of the clusters contains genes that are identified only as LOCID or other numerical designation. These are obviously the subject of ongoing curation and in some cases LOCID transcripts duplicate named transcripts in the same cluster. In **Table S2** we have included an update on candidate annotations from the RGD. Clearly, the co-expression information can provide additional assurance that putative orthology relationships with known mouse or human genes are likely to be correct.

### Transcripts that do not form clusters

The first step in network analysis is the generation of a pairwise correlation matrix, and for any gene of interest one can immediately identify others with the most similar expression patterns. By lowering the inclusion threshold (*r* value) it is possible to include a larger proportion of transcripts, but the associations may become less informative biologically. Genes with unique expression profiles across the samples will not correlate with any other and therefore will not be included in the network graph. In many cases, the unique expression profile of a gene of interest arises because the gene product is “multi-tasking” in different locations. **Figure 2** shows the individual profiles of selected examples discussed below.

**Figure 2.**
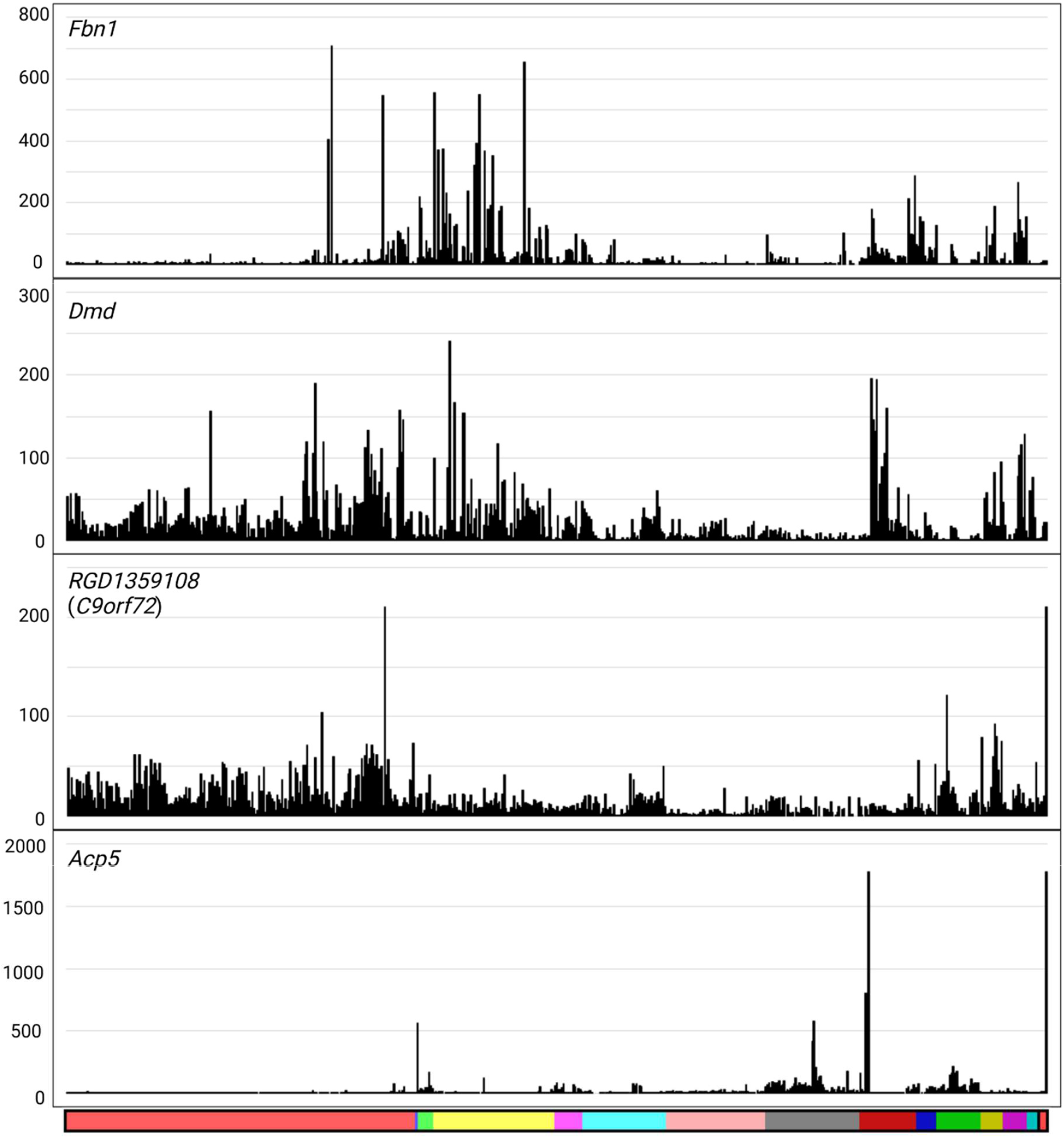
Gene expression profiles for genes which did not fall within a cluster. Y axis shows the expression level in transcripts per million (TPM). X axis shows the organ system, coloured as in **Table S2**. Reading from left to right: light red, nervous system; blue, auditory system; light green, respiratory system; yellow, cardiovascular system; pink, digestive system; turquoise, endocrine system; salmon, liver; grey, renal system; dark red, skeletomuscular system; dark blue, integumentary system; dark green, immune system; olive, male reproductive system; dark pink, female reproductive system; dark turquoise, primordia/early development; black, whole body (embryo); red, mixed tissues.

Mutations in *FBN1*, encoding the extracellular matrix protein fibrillin-1, are associated with Marfan syndrome which has complex impacts on musculoskeletal development, adiposity, vascular function and the eye. Distinct 3’ truncation mutations are associated with a neonatal progeroid lipodystrophy syndrome (54). Consistent with these phenotypes, *Fbn1* mRNA is highly-expressed uniquely in the rat eye, aorta and cardiovascular tissues and cartilage/tendons and to a lesser extent in fibroblasts and adipose. There is also moderate expression in spinal cord and dorsal root ganglia, lung and testis. Dural ectasia, enlargement of the neural canal, is a common feature of Marfan syndrome (55). Expression in the lung may underlie the pulmonary emphysema observed in mouse models of fibrillinopathy (56) patients with Marfan syndrome frequently show apical blebs in the lung and are prone to pneumothorax (collapsed lung).

The gene encoding dystrophin (*DMD*) associated in humans with mutations causing Duchenne muscular dystrophy, is also not clustered. As expected, it is expressed in rat cardiac, skeletal and uterine muscle, but is also expressed in multiple brain regions at similar levels. This expression may be related to the neuropsychiatric impacts of the disease in both affected individuals and mouse models (57). In this case, FANTOM5 data indicate that *DMD* has at least two independent promoters (6).

RGD1359108 has not been annotated on RGD, but on Ensembl it is a clear 1:1 ortholog of human *C9orf72*, associated with amyotrophic lateral sclerosis and frontal temporal dementia. O’Rourke et al. (58) reported that loss of function mutation in this gene in mice did not produce motor neuron dysfunction, but did lead to macrophage dysfunction, splenomegaly and lymphadenopathy. In rat, *C9orf72* is expressed widely in all CNS-associated tissues, most highly in spinal cord, but not enriched in any isolated CNS cell population. Outside the CNS it is most highly-expressed in stimulated macrophages and in testis.

A significant cohort of transcripts is excluded from co-expression clusters because they have alternative promoters, each with a distinct expression profile. One such gene is *Acp5*, encoding the widely-used osteoclast (OCL) marker, tartrate-resistant acid phosphatase. *Acp5* forms part of a small cluster (Cluster 179, 10 nodes) that is most highly-expressed in the femoral diaphysis, and includes another OCL marker *Ctsk*, osteoblast-associated transcripts (*Bglap, Dmp1* and *Sp7*) and *Ifitm5*, mutated in a human bone-related genetic disease, osteogenesis imperfecta type V. It is surprising that so few transcripts are stringently associated with OCL; another small cluster (Cluster 174, 11 nodes) that contains *Dcstamp, Ocstamp* (*Zfp334)* and *Mmp9*, is enriched in the diaphysis sample but more widely-expressed. Expression of *Acp5* in OCL in mice is initiated from an OCL-specific promoter (59). Aside from its function as a lysosomal enzyme in bone resorption, secreted ACP5 can function as a neutral ATPase and a growth factor for adipocytes (60,61). *Acp5* mRNA is expressed, albeit a lower levels than in bone, in rat adipose, lung (where it is expressed highly by alveolar macrophages), small and large intestine, kidney and spleen as well as isolated macrophages.

### The transcriptome of the rat liver

The downloaded datasets included around 1900 individual RNA-seq libraries of liver, including whole liver from various ages, sexes, inbred and outbred rat strains, disease models, liver slice cultures and isolated cells. In principle, clustering of such diverse data could identify sets of co-expressed transcripts that are associated with cell-types, locations or disease processes that are hidden in the averaged data of the complete sample set. To test that view, we clustered the entire liver-related dataset without averaging the replicates. As in the main atlas, the correlation threshold was chosen empirically at 0.75. The cluster list and the average profile of transcripts in each cluster is provided in **Table S3** and informative clusters are summarised in **Table 2**.

**Table 2.**
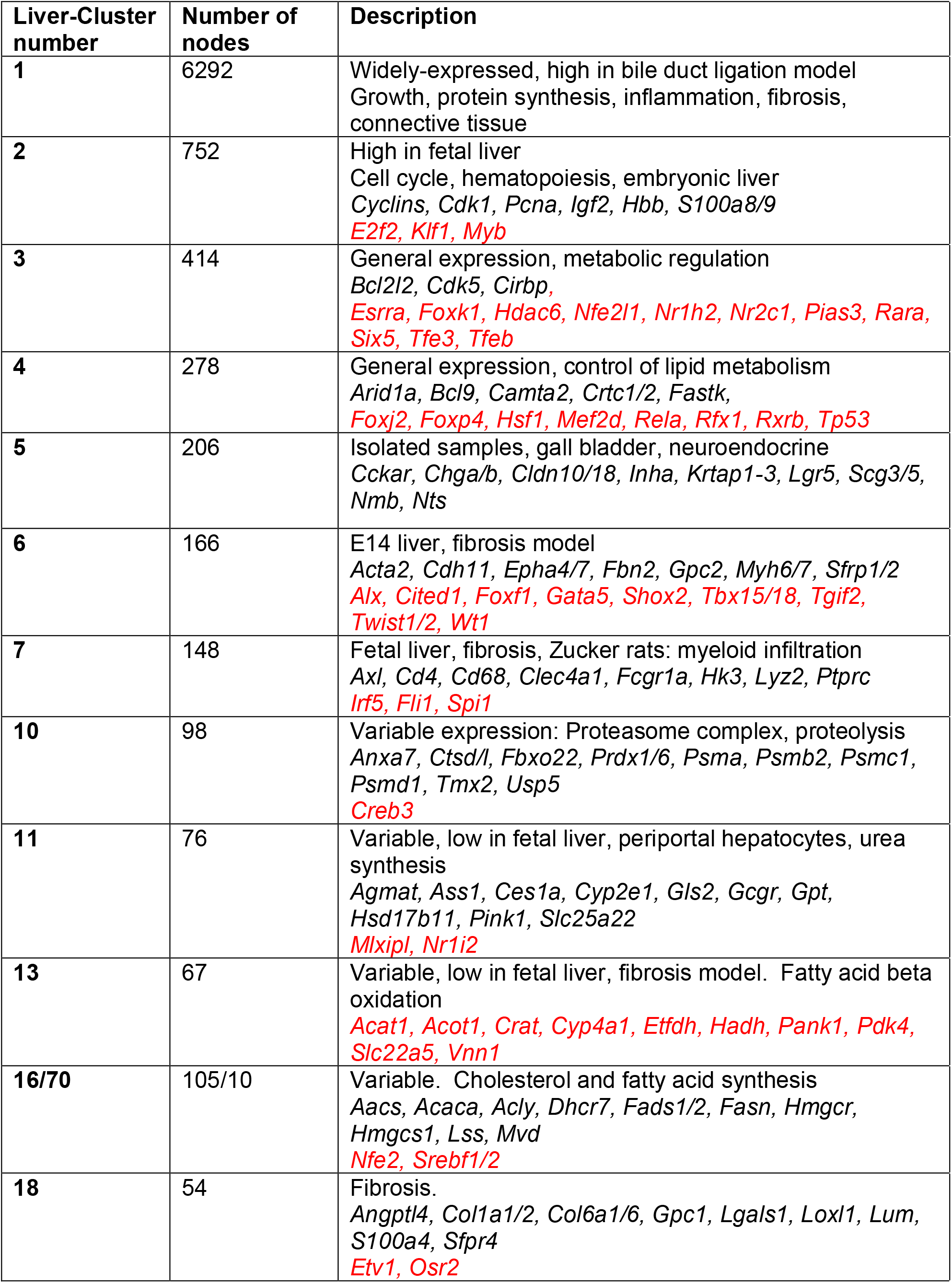
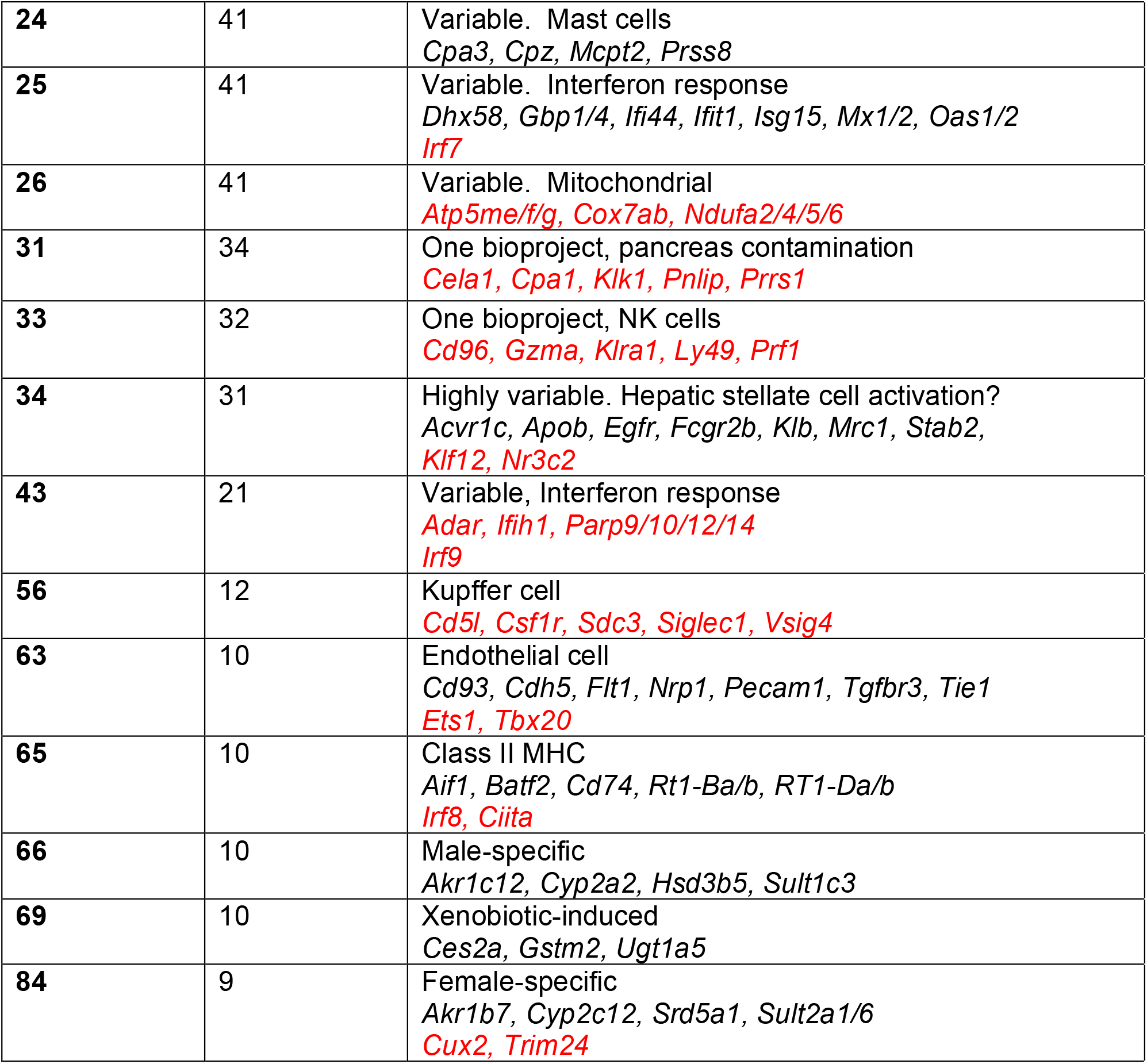
Gene expression clusters from rat liver. Clusters were generated at *r* ≥ 0.75 and MCL inflation value 1.7. Full dataset is provided in Table S3. Transcription factors are highlighted in red.

It is immediately evident that not all of the samples are pure liver. Liver-Cluster 31 contains a set of pancreas-specific genes, including *Cpa1*, that overlaps with Cluster 64 in the main atlas. This cluster arises because of random contamination with pancreatic tissue of liver samples in the large bodymap project (32). Liver-Cluster 73 contains transcripts encoding all of the major secretory products of pancreatic islets (e.g. *Ins1, Gcg*). This cluster was detected only in liver from a study of enforced activity and sleep deprivation (62). It is not clear from the paper how these samples could have been selectively contaminated with islet mRNA unless they are mislabelled. Liver-Cluster 5 is detected in a rather random subset of samples from multiple BioProjects likely also indicating contamination. It includes the progenitor marker, *Lgr5*, but also various adhesion molecules (*Cldn10/18*) and neuroendocrine markers (*Chga/b*). There is little evidence of expression of these genes in normal liver in other species, and at least some of the genes (e.g. *Cckar, Cldn10/18*) are highly-expressed in pancreas and/or stomach (e.g. see http://biogps.org). Liver-Cluster 21 is detected in a single sample, and contains smooth muscle-associated transcripts (*Actg2, Tpm2*).

The disadvantage of analysing a single tissue is that most transcripts do not vary greatly between datasets. In one sense, this provides a quality control for the efficacy of the random sampling approach we have used. In this dataset, the largest cluster by far (Liver-Cluster 1) is relatively consistent with the exception of increased detection in all samples from a BioProject that profiled liver slices from a bile duct ligation model, cultured for 48 hrs *in vitro* and treated with various agents (63). It is not clear why this gene set would be expanded in that cellular system. Liver-Cluster 1 includes many transcripts expressed constitutively by hepatocytes. The most abundant hepatocyte-specific transcript encoding albumin (*Alb*) is not strictly correlated with any other transcript presumably reflecting its specific regulation (64). Liver-Cluster 1 also contains transcripts encoding markers of hepatic stellate cells (e.g *Pdgfra/b*) and the corresponding growth factors (*Pdgfa/b/d*) as well as more general mesenchyme markers (e.g *Vim*) and markers of cholangiocytes (e.g. *Krt7*) suggesting that their relative abundance is not highly variable amongst the samples.

The remaining liver clusters analyse differential development and activation states that distinguish the samples and BioProjects. These clusters are informative and consistent with prior knowledge. Liver-Cluster 2 is expressed specifically in embryonic liver and is a complex mix of transcripts reflecting both differentiation of hepatocytes and the function of the liver as a hematopoietic organ. Accordingly, it contains the cell cycle genes, the fetal growth factor *Igf2*, and markers of erythroid (e.g. *Hbb*) and myeloid (*S100a8/a9*) hematopoietic lineages. Liver-Clusters 3 and 4 are both expressed in almost all liver samples and the level of expression is not highly variable. Expression of each of the smaller clusters is much more variable between samples and BioProjects and known genes within those clusters indicate an association with specific cell types and processes as summarised in **Table 2** and discussed below.

One signature that was no detected is that of the specialised centrilobular population that is adapted to clear ammonia generated by the urea cycle. In mice, the rate-limiting enzyme, glutamate ammonia lyase (aka glutamine synthetase, *Glul*) is expressed exclusively in a band of cells surrounding the central vein. Liver-specific deletion of *Glul* leads to pathological hyper-ammonemia (65). In mice, this population of cells co-expressed *Rhgb* (encoding an ammonia transporter) and ornithine aminotransferase (*Oat*) and was enriched for a number of *Cyp* genes (e.g *Cyp2e1, Cyp1a2*). However, in the diverse rat liver dataset, there was only marginal correlation with other centrilobular-enriched transcripts.

### The transcriptome of central nervous, renal, musculoskeletal and cardiovascular systems

Each of these systems also contributes hundreds of RNA-seq datasets including isolated cells and specific regions or structures. To examine further the utility of these large datasets for the analysis of cell-type and process-specific signatures, the data from each of these biological systems was clustered separately in **Table S4** (nervous), **Table S5** (renal), **Table S6** (cardiovascular) and **Table S7**(musculoskeletal). The clusters are annotated in the Tables and to avoid confusion with multiple Cluster numbers, each system is discussed separately in Supplementary Text. Broadly-speaking, as in the liver, network analysis of individual organ systems enables a more fine-grained extraction of cell-type, region and process-specific expression signatures.

### The transcriptome of rat macrophages

The transcriptome of rat macrophages has been analysed previously based upon microarrays (66) and the RNA-seq data included here (67). Macrophages adapt to perform specific functions in specific tissues (20). Cluster 21, which includes *Csf1r*, is most highly-expressed in brain and brain-derived cells and includes transcripts that are enriched in microglia compared to macrophages from other tissues (e.g. *P2ry12*). Around 2/3 of these transcripts are contained within a set of 119 transcripts depleted in all brain regions of *Csf1r*-knockout rats (68). Cluster 47 contains transcripts that may be shared with microglia (e.g. *Itgam*, encoding CD11b) but are common to monocytes and many tissue macrophage populations. Cell surface markers of other macrophage populations cluster idiosyncratically, indirectly supporting tissue macrophage heterogeneity; *Clec4f*, the Kupffer cell marker is within the liver cluster, *Vsig4* and *Marco* (Cluster 1239), *Clec10a, Mrc1* (CD206), and *Stab1* (Cluster 168), *Lyve1* and *Timd4* (Cluster 79), *Adgre1* and *Clec4a1/3* (Cluster 286) are correlated with each other while others (e.g. *Cd163, Tnfrsf11a, Siglec1*) do not cluster at all at this threshold because each has a unique pattern of expression in tissue macrophages. **Figure 3** shows the profiles of *Csf1r, Adgre1, Cd163, Vsig4* and *Mrc1* in the averaged data.

**Figure 3.**
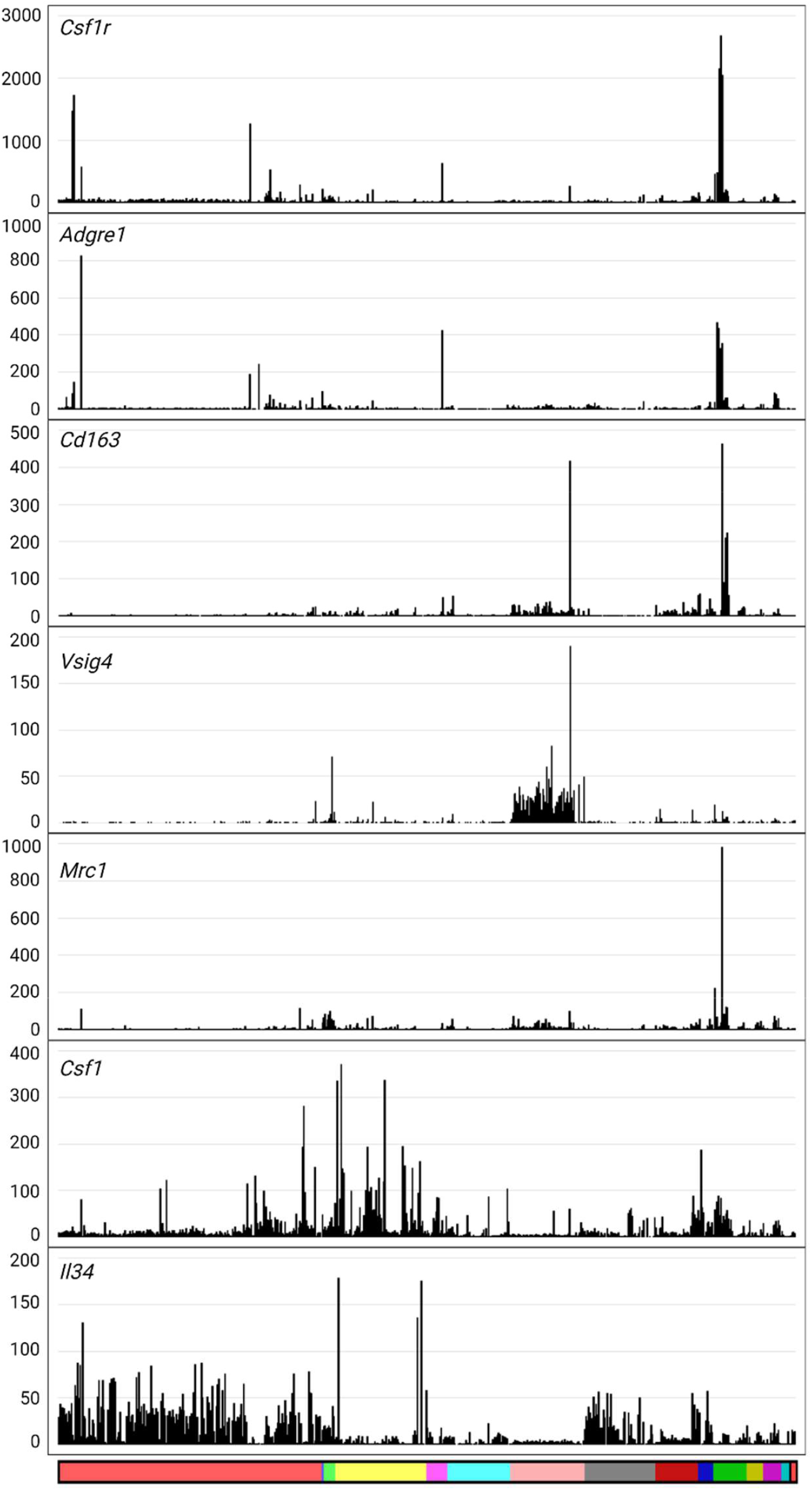
Gene expression profiles for macrophage-related genes. Y axis shows the expression level in transcripts per million (TPM). X axis shows the organ system, coloured as in **Table S2**. Reading from left to right: light red, nervous system; blue, auditory system; light green, respiratory system; yellow, cardiovascular system; pink, digestive system; turquoise, endocrine system; salmon, liver; grey, renal system; dark red, skeletomuscular system; dark blue, integumentary system; dark green, immune system; olive, male reproductive system; dark pink, female reproductive system; dark turquoise, primordia/early development; black, whole body (embryo); red, mixed tissues.

The network analysis of such a diverse set of cells and tissues also dissociates known macrophage transcriptional regulators (e.g. *Spi1, Spic, Nr1h3, Mafb, Irf8, Cebpa/b, Tfec*) (20) from macrophage expression clusters because none of these regulators is entirely macrophage-restricted. For example, transcription factor SPIC in mice is required for splenic red pulp macrophage and splenic iron homeostasis (69). In the rat, *Spic* mRNA is most highly-expressed in spleen as expected, but also detected in ES cells and germ cells. Macrophage differentiation and adaptation likely involves combinatorial interactions amongst multiple transcription factors as exemplified by the complex regulation of the transcription of the *Csf1r* gene (70).

Whereas macrophages express a diversity of endocytic receptors, there is not a corresponding large cluster of transcripts encoding endosome-lysosome components including the vacuolar ATPase (ATP6v) subunits and lysosomal hydrolases. Transcripts encoding endosome-associated CD68 and GPNMB proteins are co-expressed with *Ctsb* and *Ctsd*. Although CD68 is often used as a macrophage marker, it is clearly not macrophage restricted. Most transcripts encoding lysosomal acid hydrolases (e.g. *Acp1, Lipa*) are widely-expressed and each varies independently.

*Csf1r* is strongly correlated with other macrophage-specific markers in Cluster 21, consistent with strong evidence that expression is entirely restricted to the macrophage lineage in rats as it is in mice (71). It is also detected at relatively high levels in all tissues (around 5-10% of the level in isolated macrophages) consistent with the abundance of tissue macrophages detectable with a *Csf1r* reporter transgene (71) and with a study of tissue development in mice (72). However, expression was also detected in many isolated primary cell samples that are not meant to contain macrophages. For example, BioProjects PRJNA556360 and PRJNA552875 contain RNA-seq data derived from oligodendrocyte progenitors purified using the A2B5 marker but this population has *Csf1r* expression at similar levels to purified macrophages. Another BioProject, PRJNA355082, describes expression profiling of isolated astrocytes, but this dataset also has a similar level of *Csf1r* mRNA to pure macrophages. Other datasets from various ganglion cell populations, neuronal progenitor cells, cardiac fibroblasts and cardiomyocytes and hepatic stellate cells are clearly highly-enriched in *Csf1r* and other macrophage-associated transcripts.

CSF1R has two ligands, CSF1 and IL34. In mice and rats, mutation of the *Csf1* gene leads to a global reduction in many tissue macrophage populations, whereas mutation of *Il34* in mice leads to selective reduction of microglia and Langerhans cells. Based upon the difference in phenotype between *Csf1* and *Csf1r* mutations in rats, we speculated that *Il34* could be more widely-expressed and functional in rat macrophage homeostasis compared to mouse (67). Neither growth factor forms part of a cluster. **Figure 3** also shows the profiles of *Csf1* and *Il34*. As expected, *Csf1* mRNA is widely-expressed and enriched in isolated mesenchymal cells. *Il34* is expressed in all brain regions and isolated cells at similar levels and also in skin. However, by contrast to mouse, *Il34* is expressed at similar levels in many other tissues, notably aorta, adipose, kidney, lung and testis.

The tissue-specific analysis in **Tables S4, S5, S6 and S7** enables the extraction of macrophage-specific signatures from resident populations that have not been isolated and characterised previously. For example, in the cardiovascular analysis, a cluster of 184 transcripts containing *Csf1r* as well as a smaller cluster containing *Adgre1* extracts a signature of cardiac resident macrophages distinct from blood leukocytes which form a separate cluster (Supplementary On-line text).

## DISCUSSION

### Overview

The extraction and normalisation of published RNA-seq data has enabled the generation of a comprehensive rat expression atlas that samples transcriptional diversity on a comparable scale to the FANTOM5 data for human and mouse (6) and massively extends the *Bodymap* generated from 11 rat tissues (32). The user-friendly display at www.biogps.org/ratatlas enables a gene-specific query to visualise the expression of any gene of interest across the full dataset and use of the Correlation function allows the identification transcripts with similar expression profiles. Biogps also hosts large expression datasets for mouse, human, sheep and pig for comparative analysis. The validity of the down-sampling normalisation, and the utility and information content of the atlas has been exemplified by gene-centred network analysis (GCNA) of the averaged core dataset. The primary data is available for download by users in a form that enables local regeneration of the networks and addition of user-generated datasets. By comparison to rat, there are orders of magnitude more total RNA-seq datasets from mouse and human cells and tissues in public repositories. We previously identified and analysed 470 RNA-seq datasets from mouse resident tissue macrophages alone, excluding data from cells stimulated *in vitro* or in disease models (20). The approach we have used in extensible to even larger datasets in mouse and human.

### Analysis of liver-specific transcriptional network

The assembled dataset includes multiple BioProjects and thousands of RNA-seq datasets related to the liver, central nervous system, heart and cardiovascular system and kidney. Each has been analysed independently to identify signatures of individual cell types and processes (**Tables S3-S7**). To illustrate the ability of network analysis to extract biologically informative expression signatures, we analysed the liver data (**Table S3**) in greater detail and considered other tissue-specific analysis in Supplementary On-Line text.

Liver gene expression is regulated in response to numerous physiological stimuli and chronic disease processes including fatty liver disease. Aside from hepatic parenchymal cells, the liver contains several non-parenchymal populations. To identify co-regulated clusters within the liver transcriptome we analysed the liver samples separately using the same GCN approach used for the overall atlas. The liver is the major source of plasma protein and performs many functions in energy homeostasis, lipid and protein synthesis, biotransformation of xenobiotics and endogenous by-products. The function of the liver depends on its structure, which comprises small units called lobules each composed of concentric layers of hepatocytes expanding from the central vein toward the periportal vein. The metabolic function of hepatocytes varies along the periportal–central axis, a phenomenon referred to as metabolic zonation (73). In principle, if there was significant heterogeneity in metabolic state or development amongst the liver samples, a gene-to-gene clustering might reveal sets of genes associated with portal versus centrilobular regions of liver lobules. Halpern *et al*. (74) performed single cell RNA-seq analysis of mouse hepatocyte diversity and concluded that zonation impacts as many as 50% of transcripts. However, this analysis was limited to 8 week old fasted male C57Bl/6 mice and does not necessarily capture coordinated regulation of the metabolic domains including diurnal oscillations and response to feeding (75). Broadly-speaking, the single cell analysis indicated a periportal bias for major secretory products of hepatocytes and a pericentral concentration of expression of genes involved in xenobiotic metabolism.

Network analysis revealed a large co-regulated cluster (Liver-Cluster 11) that includes *Gls2*, an archetypal periportal marker in mice, other enzymes and transporters associated with the urea cycle (*Ass1, Acy3, Agmat, Cbs, Gpt, Slc25a22, Nags*) and the glucagon receptor, *Gcgr*. Cheng *et al*. showed that glucagon is a regulator of zonation in mouse liver, in that glucagon deficiency led to reduced expression of periportal-enriched transcripts (76). There are candidate transcriptional regulators within this cluster with known functions in hepatic transcriptional regulation; the xenobiotic sensor *Nr1i2* and the glucose-sensing transcription factor *Mlzipl* (77,78). A smaller Liver-Cluster 88 contains additional key enzymes of urea synthesis, *Arg1, Cps1, Gpt2* as well as the amino acid transporter, *Slc38a4*.

The analysis does not reveal a corresponding pericentral expression cluster. *Glul*, which appears strictly-restricted to a single layer of cells surrounding the central vein in mice, rats and humans (73) showed limited heterogeneity amongst the liver datasets and did not form part of this cluster. This suggests that *Glul* is not highly-regulated whereas other centrilobular-enriched transcripts alter their expression in response to external stimulus. Another putative landmark pericentral gene, *Cyp2e1*, is actually part of Liver-Cluster 11, redistributed in at least some of the experimental models sampled herein, as observed in a model of paracetamol exposure that forms part of this dataset. Other transcripts that are biased to centrilobular also form separate clusters because of their independent regulation in response to stimulation. For example, *Cyp1a2* was identified as a pericentral marker (73). Liver-Cluster 54 is elevated in a dataset from a BioProject studying the effects of 2,3,7,8-Tetrachlorodibenzo-p-dioxin (TCDD), a potent aryl hydrocarbon receptor (AhR). It includes the detoxifying enzymes *Cyp1a1, Cyp1a2 and Cyp1b1*, the AHR repressor gene *(Ahrr*) and transcription factor *Cdx2*, a known AHR target gene (79). A distinct set of xenobiotic metabolising genes, *Ces2a, Gstm2* and *Ugt1a5* is coregulated in Liver-Cluster 69, and *Ephx1* and *Gsta2,4,5, m1* are co-regulated in Liver-Cluster 146. The proteasome subunit, *Psmd4* was also pericentral in mice (74) but it is found in Liver-Cluster 10 stringently co-regulated as one might expect with numerous other components of the proteasome complex. Liver-Cluster 10 contains the transcription factor *Creb3*, and likely reflects the activation of the Golgi stress response in a subset of samples or BioProjects (80).

The regulation of lipid metabolism is of particular interest given the current epidemic of non-alcoholic fatty liver disease. There is some evidence of zonation of fatty acid metabolism in the liver; fatty acid β oxidation being enriched in periportal and lipogenesis in pericentral hepatocytes (74) but these pathways are independently regulated in this dataset. Liver-Cluster 13 is highly-enriched for genes involved in lipolysis and fatty acid β oxidation. It overlaps the smaller cluster in the full atlas (Cluster 101) but includes many additional genes that have tissue-specific enrichment (e.g. *Acot7* in CNS). Conversely, Liver-Clusters 16 and 70 comprise enzymes of cholesterol and fatty acid synthesis and the known transcriptional regulators, *Nfe2* and *Srebf1/2*. Liver-Cluster 26 contain multiple genes involved more generally in mitochondrial oxidative phosphorylation including multiple genes encoding NADH-ubiquinone oxidoreductase (NDUF) subunits. We are not aware of any heterogeneity in mitochondrial distribution in the liver.

The various metabolic and inflammatory disease models, with distinct effects on non-parenchymal cells, enable deconvolution of signatures of specific cell types and disease processes. Liver-Cluster 6, which includes the classical fibrosis marker, *Acta2* (smooth muscle alpha actin/ SMA) is elevated in fibrosis models, but highest in E14 liver, which may indicate that myofibroblast activation in fibrosis recapitulates the phenotype of embryonic mesenchyme. Liver-Cluster 18 captures transcripts associated with more advanced fibrotic disease and includes multiple collagen genes and two candidate transcriptional regulators, *Etv1* and *Osr2*. This cluster also contains the mesenchymal gene *Olfml3*, which is also expressed in microglia in the mouse (see biogps.org) and human (81) but is not associated with microglia in the rat (68). This highlights the problems with assuming that genes have similar expression patterns and functions across species.

The fibrosis-associated clusters are clearly separated from Liver-Cluster 7 which captures the phenotype of infiltrating CD45^+^ (*Ptprc*) myeloid cells in various models. Two sets of interferon-responsive transcripts including key regulators *Irf7* and *Irf9* cluster separately (Liver-Clusters 25 and 43) as do transcripts associated with expression of class II MHC (Liver-Cluster 65). These clusters are separated also from the signatures of endothelial cells (Liver-Cluster 63) and of Kupffer cells, the resident macrophages (Liver-Cluster 56). The latter cluster includes the transcript encoding the macrophage growth factor receptor, *Csf1r* and many transcripts that were also down-regulated in livers of *Csf1r-*knockout rats (82). *Clec4f*, which is expressed exclusively by Kupffer cells in mice, and is in the liver-specific cluster in the extended atlas, is in a separate cluster (Liver-Cluster 95) with the three C1q subunits (*C1qa/b/c*), *Cfp, Ctss, Pld4* and *Tifab*. There is emerging interest in the later gene, a forkhead-associated domain protein, in immune cell function and inflammation (83).

Finally, in rodents, there is a set of transcripts that is expressed in the liver in a sex-specific manner in part under the influence of growth hormone (84,85). The male and female-specific liver transcriptomes are regulated by differential expression of specific transcription factors, CUX2 and ONECUT2 in females and BCL6 in males. The majority of samples are from males, but nevertheless, Liver-Cluster 66 is excluded from female livers, and Liver-Cluster 84 contains *Cux2, Trim 24* and known female-specific transcripts.

### The relationship between network analysis and single cell RNA-seq for the definition of cell types in tissues

As in the liver, the network analysis of other major organ systems enabled robust extraction of clusters of co-regulated transcripts often including the transcription factors that regulate them. In this case, the issue of tissue-specific promoters becomes less of an issue and genes that have multiple promoters (e.g. *Mitf, Acp5*) may form part of tissue-specific networks highlighting local functions. The deconvolution of large datasets by network analysis complements single cell RNA-seq (scRNA-seq) analysis which has rapidly become a dominant approach to analysis of cellular heterogeneity. scRNA-seq is not quantitative. Typically, expression of <1000 genes is detected in each cell and even the most highly-expressed transcripts are not detected in every cell (86). The output of scRNA-seq conflates two distinct types of zero values: those where a gene is expressed but not detected by the sequencing technology (stochastic sampling) and those that reflect genuine expression heterogeneity. Whereas we can readily separate entirely unrelated cells that share few markers in scRNA-seq, such as epithelia and hematopoietic cells, the identification of numerous subpopulations within individual lineages is tenuous at best (20). A second disadvantage of analysis of isolated cells by scRNA-seq or total RNA-seq is that cells are inevitably activated during isolation and single cells can have attached remnants of other cells that contribute RNA (20).

Suo et al. [87] described computational analysis of mouse cell atlas to identify 202 regulons whose activities are highly variable across different cell types and predicted a small set of essential regulators for each major cell type in mouse. We have achieved the same outcome for the rat without the use of scRNA-seq. The advantage of network deconvolution as performed here is that one can explore a much wider diversity of states than can be contemplated with scRNA-seq and identify more robust co-regulatory modules. Any proposed pair of markers of a specific cell population defined by scRNA-seq should be strongly correlated with each other if both are detectable in whole tissue. The prediction was tested in a meta-analysis of mouse tissue macrophage populations which failed to support the existence of a specialised macrophage subset defined from scRNA-seq data by reciprocal expression of *Lyve1* and *Mrc1* (20). Herein the detailed analysis of the liver data indicates that zonation of the liver is dynamic and individual pathways are regulated to a large extent independently of each other. So, the definition of subpopulations of hepatocytes is state-dependent. The discussion of other systems in Supplementary On-line text casts doubt on the fine-grained definition of subsets of tissue-specific parenchymal/epithelial cells and more generic glial cells, fibroblasts, endothelial cells, parenchymal cells and macrophages in many published scRNA-seq analyses. Network analysis reveals regulons that may, or may not, be restricted to a defined cell population, but which are clearly linked to function. In that respect one might reasonably question the value of defining cell types as an approach to understanding biology.

## Author contributions

SJB developed the informatics pipeline and generated the primary expression data. KMS performed the network and enrichment analyses and manual annotation of metadata. CW developed BioGPS and established the BioGPS viewer of the atlas. DAH wrote the initial manuscript, reviewed all genes and clusters and contributed to informatic analysis. SJB and KMS contributed to manuscript editing.

## Supplementary On-Line Text

### Cluster analysis of nervous system samples

The largest collection of individual RNA-seq datasets in the atlas is related to central and peripheral nervous tissues and includes 1855 samples. **Table S4** lists all of the samples and the set of Clusters identified by gene-centred network analysis. Brain region-specific analysis in juvenile rats has been reported previously (1) and here we will not attempt a detailed annotation of every cluster. There are obvious clusters of neuronal cell types enriched for specific neurotransmitter receptors or functions and specific transcription factors. For example, Cluster 4 is enriched in dorsal root ganglia (DRG), and contains specific transcription factors, *Drgx* and *Isl2*. The smaller Cluster 65 is even more DRG-restricted and contains the nociceptor marker *Ntrk3* (TRKA)(2), pain-associated receptors (*Prokr1/2*) and transcription factors *Hmx1, Isl1, Pou4f1* and *Prdm12*. Cluster 23 contains *Kit, Slc1a1, Gria1/2* and *Htr1a* and multiple voltage-gated potassium channels, Cluster 25 contains *Ntrk3* and *Grm3,5,7*, Cluster 46 contains transcripts expressed in cerebellum and Cluster 55 clearly has a signature of dopaminergic neurons including transcripts encoding synthetic enzymes (*Dbh, Ddc, Maoa, Th*). Finally, the small Cluster 295 contains multiple neuron-specific transcription factors (*Bcl11a, Fezf2, Foxg1, Lhx2, Neurod1* and *Tbr1*) that have each been implicated in aspects of axonal guidance (3).

**Figure S1.**
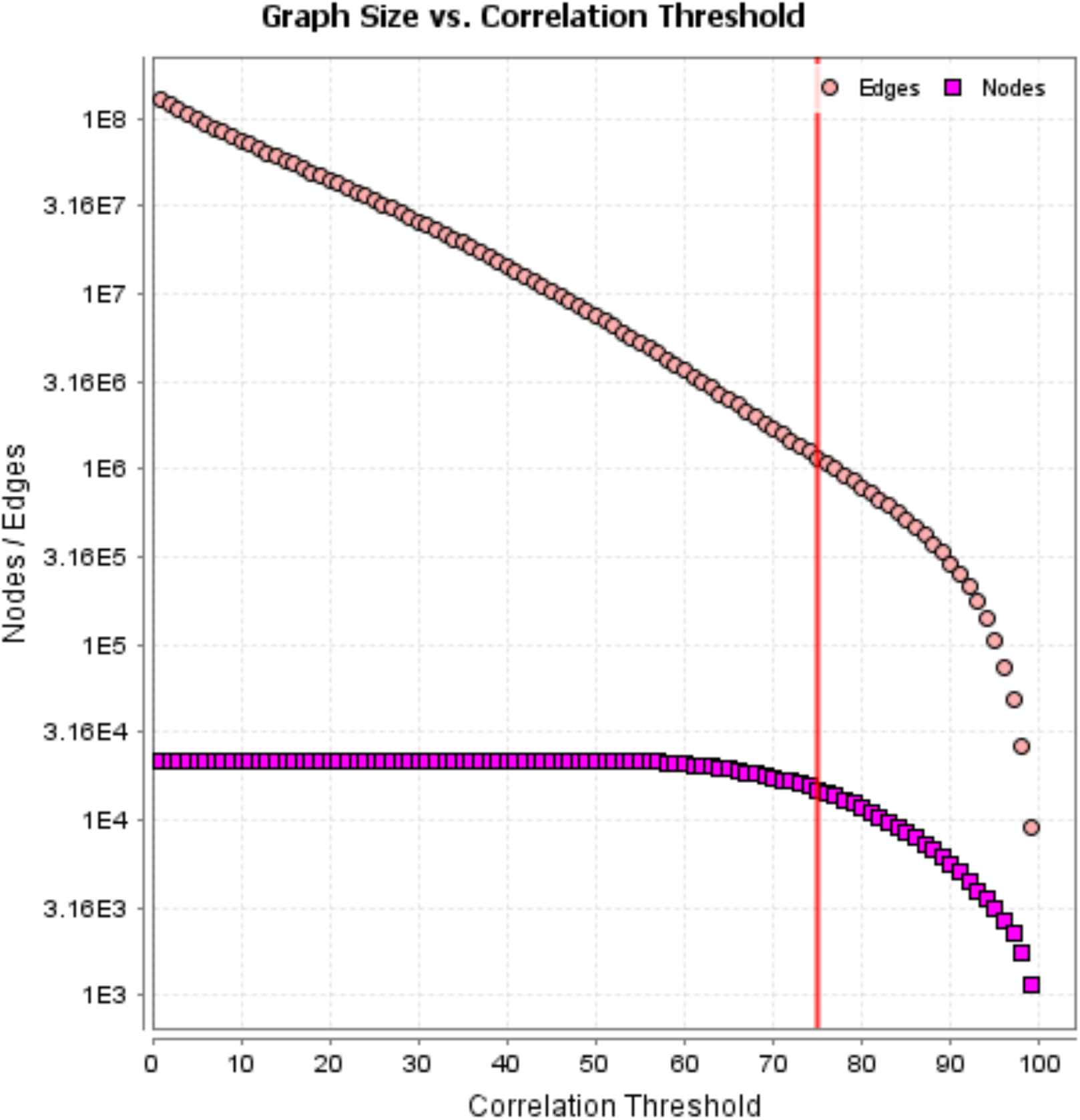

Cluster 5 is microglia-related, whilst the separate small Cluster 150 contains markers of brain-associated macrophages (e.g. *Mrc1*/CD206). Cluster 6 is expressed in pineal gland, Cluster 14 contains the transcripts for the structural and regulatory components of motile cilia, some of which are shared with testis in the main atlas, Cluster 15 contains smooth muscle alpha actin (*Acta2*), *Pdgrb* and various collagen genes and likely provides a signature of pericytes whereas endothelial markers (e.g. *Pecam1*) are in Cluster 150 alongside brain-associated macrophage markers. Transcripts associated with myelination (*Mag, Mbp, Mog, Plip*) are co-expressed in Cluster 36, although separated from the oligodendrocyte progenitor-specific transcription factors, *Olig1* and *Olig2*. This separation occurs because of the inclusion of an oligodendrocyte progenitor population purified using the surface marker A2B5 (4). The original report claims minimal contamination with microglia (<0.8%) but in fact these cells have the highest expression of any sample of the microglia-associated transcripts. They express *Olig2*, but also lack expression of mature oligodendrocyte markers associated with myelination (e.g. *Mog*). *Sox10*, which is also expressed in these cells and known to be involved in oligodendrocyte differentiation (4) is actually in Cluster 32, a Schwann cell-enriched cluster, alongside surface markers (*Cadm4, Fermt2, Itga7, Mcam*; (5)), multiple genes involved in their regulation and function (e.g. *Erbb2/3* (6), *Dhh, Bmp1, Matn2*, semaphorins (*Sema3a/3g*) and several laminins (7). The neurotrophic chemokine meteorin-like (*Metrnl*), also within this cluster, has not previously been attributed a function in Schwann cells.

We do not detect an astrocyte-specific cluster containing any of the conventional markers such as *Aldh1l1, Gfap, S100b, Slc1a2/3* or *Aqp4*. Mays *et al*. (8) reported scRNA-seq analysis of rat pineal gland and identified 3 distinct astrocyte populations, but close examination of their data suggests a poor correlation between the markers. Recent scRNA-seq data analyzing mouse cells harvested using an *Aldh1l1-*EGFP reporter also indicates these cells are extremely heterogeneous and each of the markers is independently-regulated (9). The use of *Aldh1a1* as an astrocyte marker is difficult to justify. The gene product has no known function in astrocytes; and is almost undetectable in rat brain or in human astrocytes (10). It is part of the liver-specific cluster in the main atlas, and studies of the mouse knockout focus on hepatic function and tumorigenesis (11). The simplest interpretation of the mouse scRNA-seq data is that the *Aldh1a1* marker is actually not astrocyte-specific. Batiuk *et al*. (12) used a different marker, ATP1B2 (also not clustered in our dataset) and scRNA-seq to isolate and identify 5 separate region-enriched astrocyte populations in mouse brain. The heterogeneity of astrocytes has been recognised for many years. For example, Waltz and Lang (13) used IHC to locate 3 putative markers (GFAP, glutamine synthase (*Glul*) and S100B) in rat hippocampus and concluded that up to 40% of astrocytes were GFAP-negative and GFAP-positive cells were selectively expanded in injury-associated gliosis. We do in fact identify a very small cluster (Cluster 359) that contains *Glul* and other markers enriched in rat astrocytes, the neurotensin 2 receptor (*Ntsr2*), *Aldoc* and *Gjp6* (14-16). This cluster supports Claudin 10 (*Cldn10*) as an additional marker. *Cldn10* is detectable in rat and mouse brain, albeit lower than in kidney. These may be the only robust astrocyte markers in the rat.

Analysis of the averaged data in the full atlas dataset revealed a cluster of transcripts enriched in neurogenic progenitors. This cluster containing the commonly-used marker, *Dcx*, is further expanded in the CNS restricted dataset. Cluster 13 includes multiple known surface markers (e.g. *Cdh4, Cd24, Cxadr, Gpr85, Lrp8)* of neurogenic cells, known and novel transcriptional regulators (*Hdac2, Hes6, Mycl, Mycn, Sox4/11/12*) and tubulin subunits (*Tuba1a, Tubb2b, Tubb5*). By extension, many other genes in this cluster likely have a function in neurogenesis and are candidate genes for involvement in human lissencephaly (absence of folds in the cerebral cortex) associated with failures of neurogenesis and neuronal migration (17).

### Cluster analysis of renal samples

The atlas dataset includes RNA-seq data from 17 separate BioProjects of the renal system (**Table S1**) including studies of isolated cells, dissected regions, diabetes, injury and disease models and effects of age, developmental stage and effects of mutations. Each of these BioProjects provides multiple replicates. The kidney data include datasets from micro-dissected renal tubules (18), an analysis that is more practical in the rat than the mouse. A subsequent study in mouse (19) proposed the existence of signatures of as many as 43 separate cell types in the total kidney RNA-seq data based upon specific markers and attempted to integrate with numerous published scRNA-seq datasets from mouse kidney. Previous efforts to deconvolute whole tissue data to extract cell-specific signatures used single cell data as a reference (20). A recent study reported eQTL analysis of microdissected human kidney samples to identify cell-type specific eQTL that in turn linked to some 200 genes regulating kidney function and blood pressure (21) and highlighted specific markers of the major cell populations within the kidney.

**Table S5** shows the set of co-expression clusters extracted from the rat renal RNA-seq data and these are summarized in the Table below. Consistent with evidence that proximal tubules contribute the bulk of mRNA, the largest cluster contains numerous known markers enriched in proximal tubules including 62 solute carriers and many transcriptional regulators known to be involved in renal development of functional regulation. Clusters 2,3,4 are associated with specific BioProjects and Cluster 5 is the cell cycle cluster, in this case elevated in a model of unilateral nephrectomy. Cluster 19 is surprising in that it contains *Alb* and *Afp* and includes an array of transcripts encoding plasma lipoproteins, complement, and clotting factors normally associated with the liver. This cluster is attributable to inclusion of one embryonic kidney sample from the developmental series and is presumably due to misidentification or contamination.

Broadly-speaking, the analysis demonstrates that it is possible to extract the signatures of all of the major cell types of the kidney and identify candidate regulators of their expression without disaggregation or isolation or the use of single cell RNA-seq. This outcome includes a clear separation of principal cells and intercalated cells from the collecting duct. Cluster 18 contains the markers of principal cells. Interestingly, the cluster also contains the peripheral neuronal marker *Ntrk1*, but no other markers of neurons. A recent study identified an *NTRK1* mutation segregating with bipolar disorder and an inherited kidney disease (22). The latter phenotype was attributed to mutation in the neighbouring *Muc1* gene, but *Muc1* is barely detectable in kidney and not part of a cell-specific cluster. Chen *et al*. (23) distinguished intercalated cells in the mouse based upon expression of KIT (*Kit*; which is grouped with its ligand, *Kitlg*, in Cluster 9). They suggested that expression of two markers, *Slc4a1* and *Slc26a4* was mutually exclusive and identified putative markers of type A and Type B intercalated cells. However, the conclusion was based upon a small number of cells and in our analysis none of these markers defines a separate cluster. One other notable feature of our analysis was the identification of a clear signature of resident kidney tissue macrophages including the receptor for the macrophage growth factor, *Csf1r*. Macrophages detected using the F4/80 marker in mice, or *Csf1r* reporters in mice and rats, are abundant in the medulla, providing an almost continuous lining of the epithelial basement membranes (24-26), but they are clearly under-represented in all published scRNA-seq datasets. In common with many other tissue macrophages, these cells express C1q subunits. As noted in the main text, a novel feature of these kidney macrophages that we have not observed elsewhere is their expression of multiple other components of the classical complement pathway and the Fc receptors, *Fcrm* and *Fcrma*. However, our analysis provides no support for CD81 as a proposed marker of resident rat renal macrophages (27)

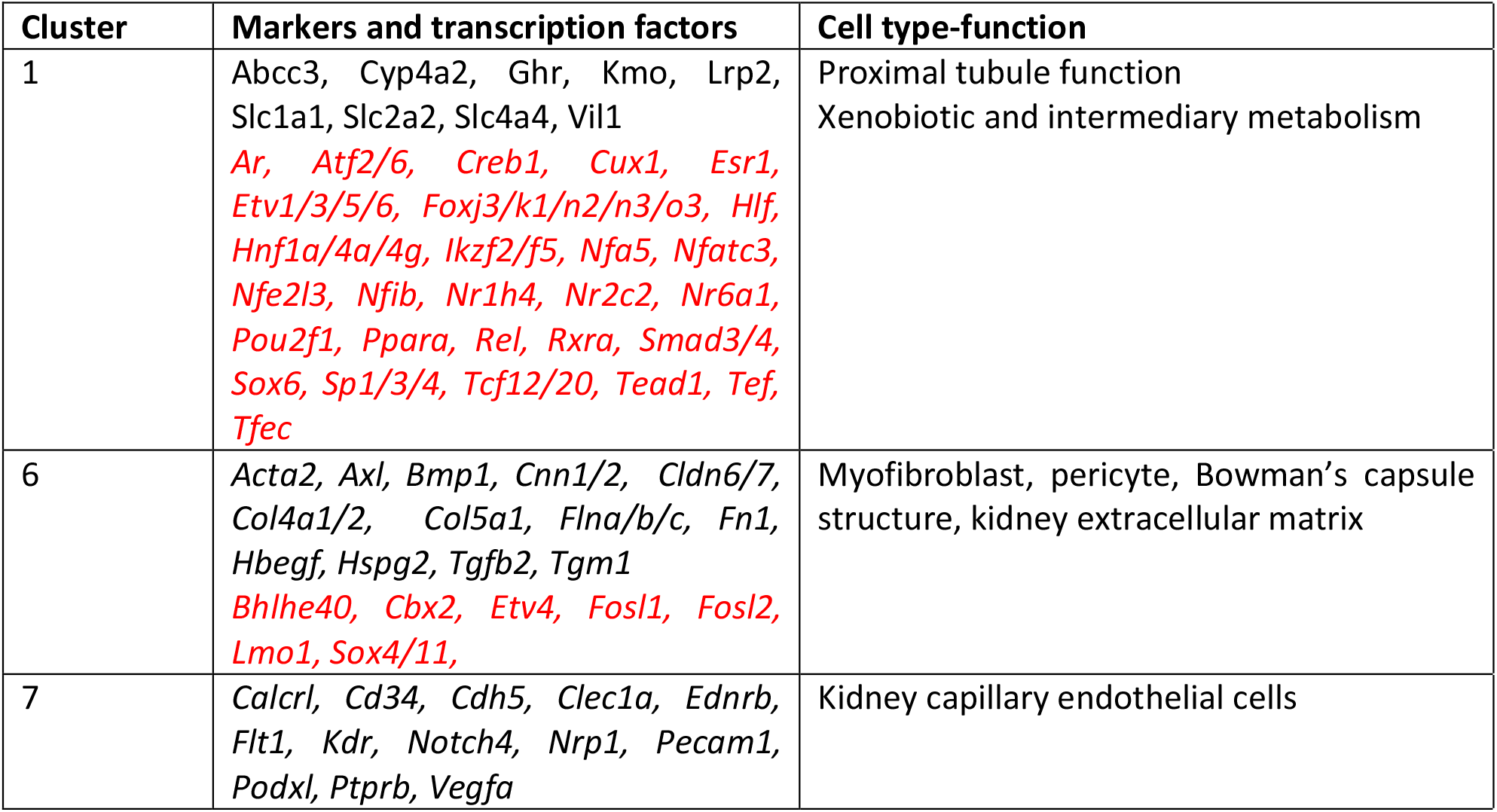
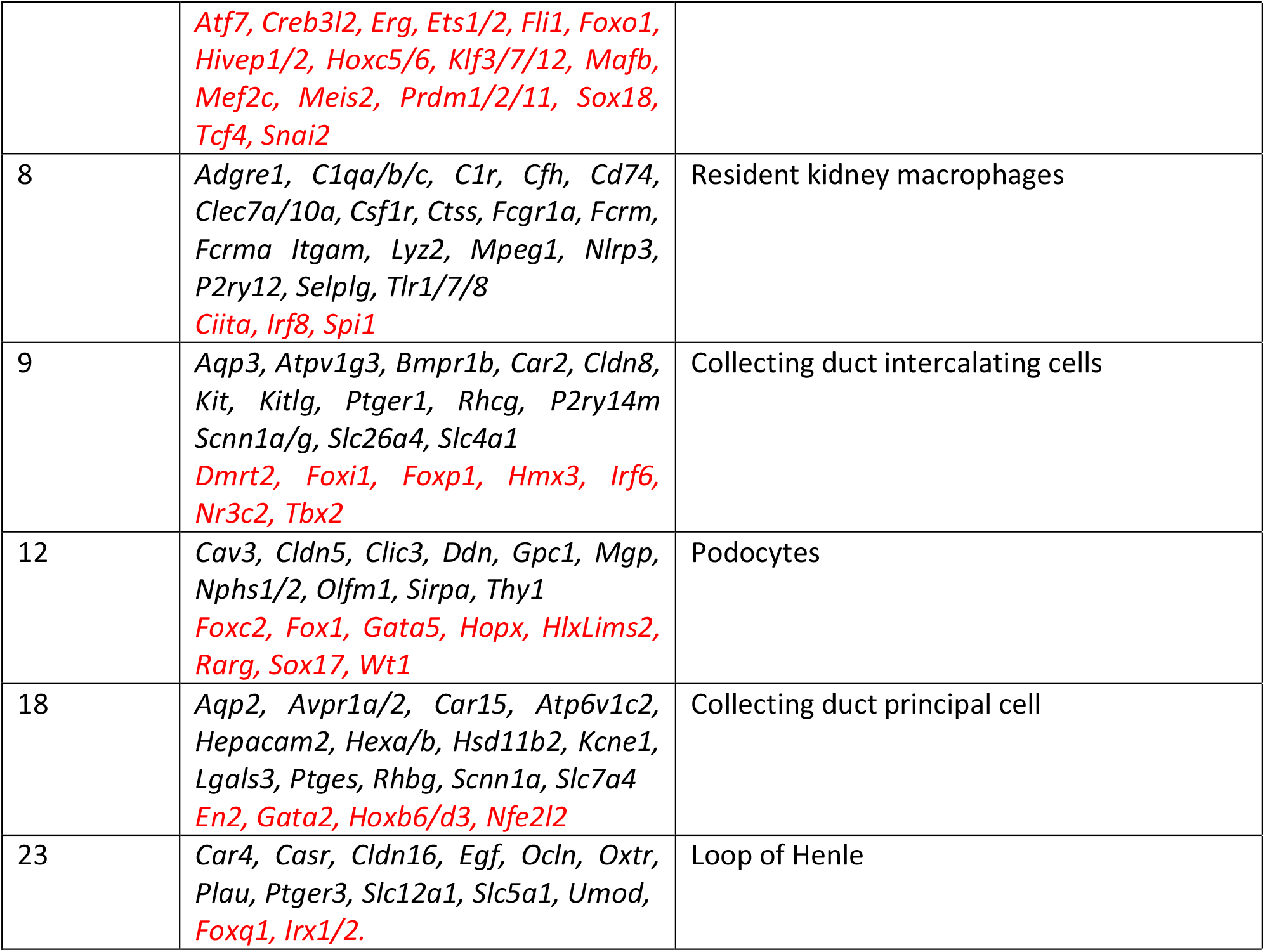

### Cluster Analysis of Cardiovascular Tissues

Cardiovascular tissues are presented by 25 BioProjects and include major vessels, intact heart, heart regions and isolated cells at different developmental stages. In common with every other organ, there have been multiple published datasets exploring cell-types in heart based upon scRNA-seq (reviewed in (28)). Each of these studies identifies numerous subpopulations of cells. An analysis of non-cardiomyocyte populations in the mouse claimed the existence of 30 distinct cell types including 8 distinct populations of macrophages (29).

**Table S6** lists the clusters identified from gene-centred network analysis of all of the individual cardiovascular-related datasets and these are summarized in the Table below. As in other datasets, there is evidence of contamination with unrelated tissues; for example Cluster 3 contains surfactant protein transcripts and likely reflects inclusion of lung tissue. Cluster 5 contains markers of B cells (*Cd19*) and T cells (*Cd3*), and likely reflects contamination with thoracic lymph nodes and Cluster 7 derives from a single sample of mesenteric artery and is likely an intestinal contaminant.

The largest cluster in this dataset with >3500 nodes is enriched in all of the isolated primary cells and is clearly associated with cell growth and proliferation. The cluster includes multiple transcriptional regulators, some of which are generic to cell cycle regulation (e.g. *Foxm1, E2f, Myc*) whilst others such as *Meis1, Runx1* and various *Smad* and *Tcf* factors (30,31) have well-defined specific functions in cardiomyocyte proliferation and development.

Cluster 4 is the major cardiomyocyte-specific cluster, and consistent with the high metabolic demand of these cells this cluster also contains multiple transcripts associated with oxidative phosphorylation. There is some evidence of independent regulation in that the large majority of components of the electron transport chain are clustered separately (Cluster 28), and the mitochondrially-encoded transcripts are also separated (Cluster 105). Cluster 17, which likely defines a distinct cardiomyocyte regulon, includes *Cav3* and components of the sarcospan complex, which can mitigate pathology in muscular dystrophy models (32). Disruption of the sarcospan complex causes cardiomyopathy in mice (33)

Broadly-speaking, the data provide little support for the extensive subset identification amongst fibroblasts, endothelial cells, pericytes, adipocytes and macrophages in published mouse and human scRNA-seq data. Each of these populations is clearly distinguished from the others but is represented by a single large cluster containing markers that are said to distinguish subpopulations in scRNA-seq data. If cell subtypes do exist, the differences between them are too subtle to enable the extraction of a signature.

Cluster 10 defines a resident cardiac macrophage population including the lineage-restricted receptor *Csf1r*. A separate Cluster 81 containing macrophage markers *Adgre1* and *Mrc1* may reflect some regional heterogeneity between the heart and aorta, which also contains a substantial macrophage population (34). We do not detect signatures of monocytes (e.g. *S100a8, Ccr2, Ly6c*) that have been reported in scRNA-seq studies. As in kidney, we suspect that disaggregation approaches provide a poor recovery of intact resident macrophages relative to recent arrivals that may be transiting through the tissue (35). The samples include genetic disease models and the power of the cluster analysis is evident in the separation of two interferon-related regulons, Cluster 6 containing *Irf9* and Cluster 23 containing *Irf7* and their respective target genes. The separation of these two interferon target cohorts was identified previously in human macrophages (36).

The cluster analysis also reveals the signature of innervation of the heart. The heart has a substantial intrinsic autonomic nervous system involved in cardiac pace-making and conduction (reviewed in (37)). This system has not been effectively profiled in scRNA-seq data, presumably because neurons are not accessible to tissue disaggregation. Cluster 18 includes the regulatory receptors *Ntrk1* and *Ngfr*, key enzymes of dopamine metabolism (*Th, Ddh*), dopamine receptor *Drd2* and other neurotransmitter receptors. Clusters 13 and 15 also contain neuronal markers. *Ntrk3*, which is associated with congenital heart disease in humans, is in Cluster 15 (38).

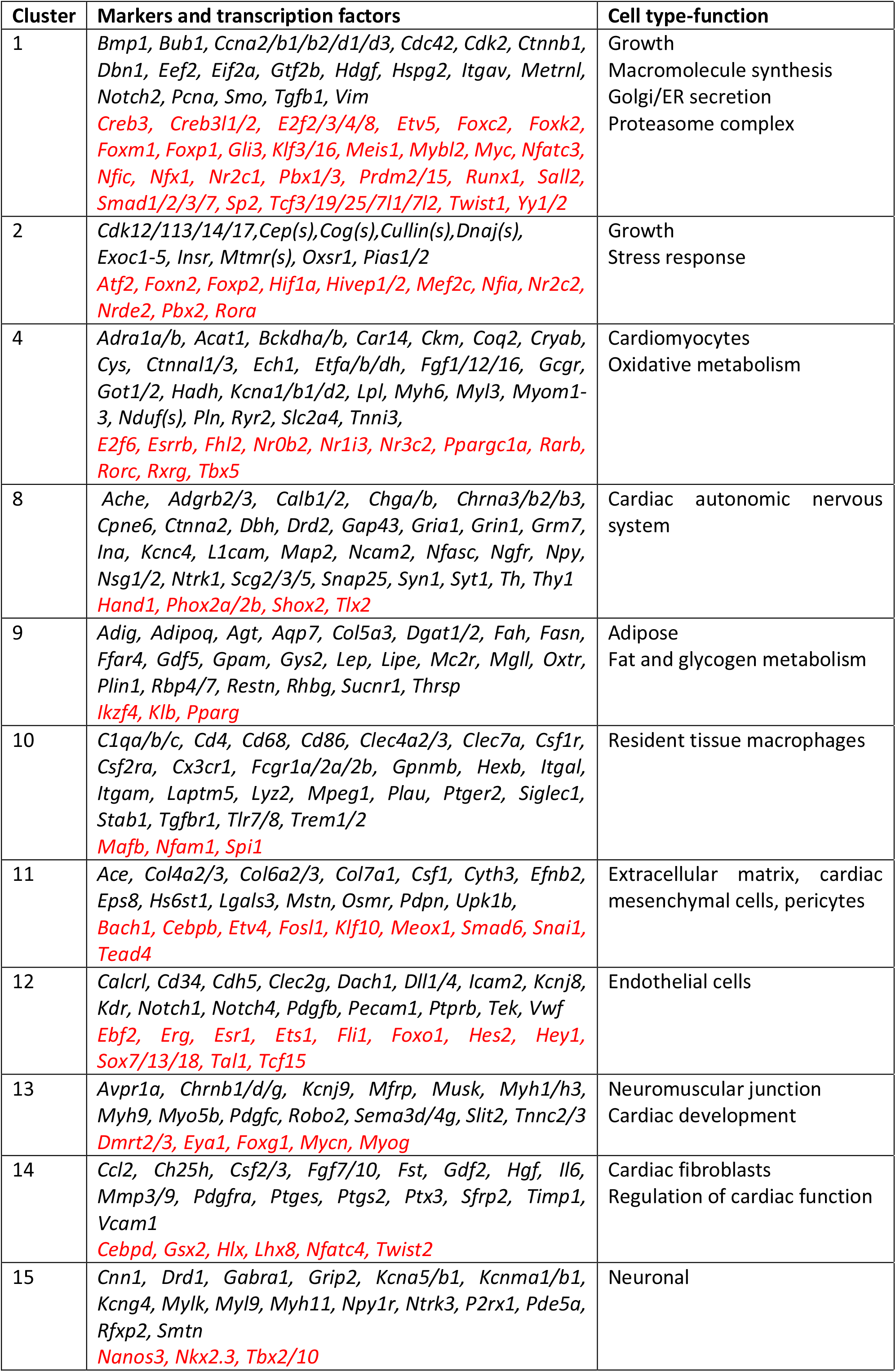
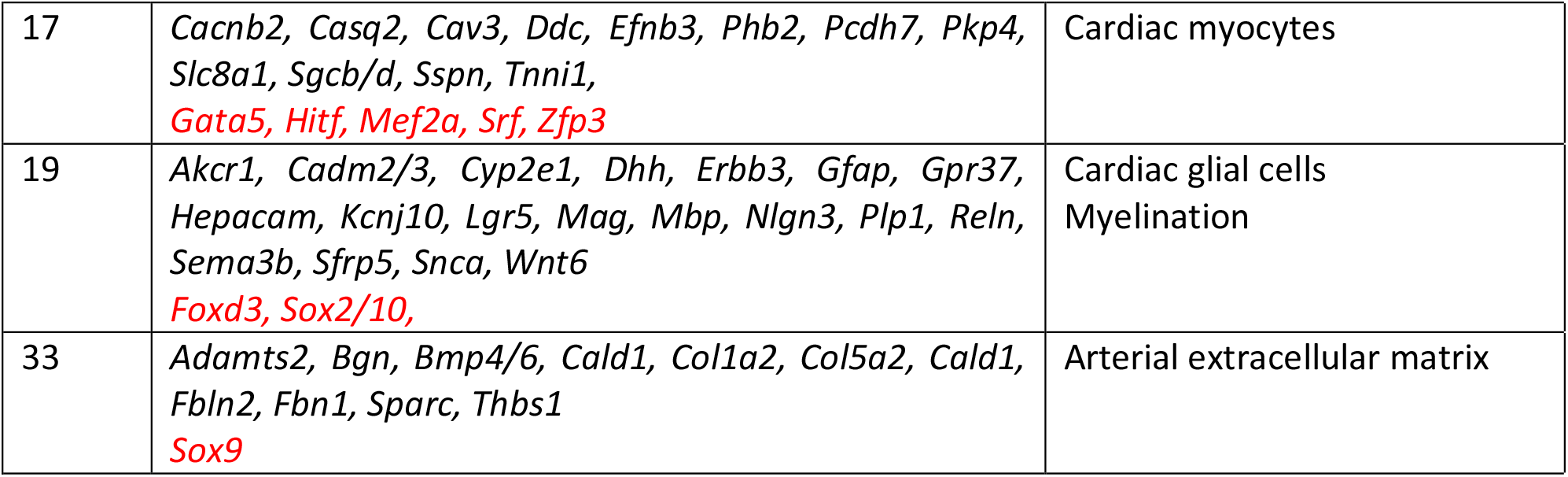

### Cluster analysis of musculoskeletal tissues

The musculoskeletal category includes samples from 33 BioProjects (Table S1), including muscle from different locations and ages as well as bone, cartilage and tendon. Unlike other groupings, the set analysed here does not include isolated cells or dissected regions or genetic disease models and accordingly the representation of some cell types is relatively homogeneous. **Table S7** contains the lists of clusters from a gene-centred network analysis of these samples. Because of the relative homogeneity of these tissues, the analysis was performed at two different MCL inflation values; clustering at an MCL inflation value of 1.7 alters the granularity but the two largest clusters remain almost unchanged when clustered at an inflation value of 2.2. For the purpose of consistency, we discuss clusters identified at MCL 2.2 used in other analyses. The largest cluster contains 4433 transcripts. Reflecting the abundance and relatively uniform distribution of interstitial macrophages in muscle and connective tissue detected with a *Csf1r* reporter transgene in both mice and rats (24,25). Cluster 1 contains *Csf1r* and many other macrophage-expressed transcripts encoding surface markers (*Adgre1, Cd14, Cd163, Cd4, Cd68, C1q, Cx3Cr1, Fcgr1, Mpeg1, Mrc1, Siglec1*) and transcription factors (*Cebpa, Irf8, Mafb, Spi1*) in common with cardiac muscle macrophages. These transcripts are separated from Cluster 27, which includes *Itgam* (Cd11b) and *Itgax* (Cd11c), generally considered markers of inflammatory macrophages in rat skeletal muscle (39). Interestingly, Cluster 1 contains the gene for the CSF1R ligand *Csf1*, and the transcript encoding the other CSF1R agonist, *Il34*, is also detected in muscle and contained within Cluster 8 with markers of adipocytes and endothelial cells and other growth factors, notably *Igf1*.

The analysis of smaller clusters reveals regulons associated with specific cell types and processes. We were interested in whether the analysis might identify components of the neuromuscular junction (NMJ) and satellite cells, which together control muscle homeostasis and regeneration. Many human genetic and acquired disease states, as well as normal ageing-related sarcopenia, impact this structure (reviewed in (40,41)). The structure and functions of the NMJ and satellite cells are tightly-linked and we anticipated that clustering would group components of both cell populations. Indeed, Cluster 14 contains transcripts encoding the cholinergic receptors of the NMJ (*Chrna1, Chrnd/e/g*) and muscle receptor tyrosine kinase (*Musk*) alongside the satellite marker *Ncam1* and myogenic determining transcription factors *Myf5, Myod1, Myog* and *Runx1*, the latter essential for satellite cell activation during muscle regeneration (42). Another transcription factor in this cluster, *Scx*, is also associated with progenitor populations albeit more commonly associated with bone and tendon (43). *Pax7* which is required for specification of satellite cells and commonly used as a marker (44) does not form part of this cluster. PAX7 protein is expressed in rat satellite cells (45) but the *Pax7* transcript is not actually detectable at >10TPM in total muscle mRNA. The other key NMJ marker, acetylcholinesterase (*Ache*) may have distinct regulation and is part of a smaller cluster (Cluster 158). That cluster includes *Sema6c*, which has been implicated in neuromuscular junction formation (PMID: 17605078).

Cluster 14 contains many novel transcripts that are known or candidate regulators or structural components but have not been widely studied. One novel member of this cluster is *Spg21*, associated with the human neuropathy Mast syndrome (hereditary spastic neuralgia). Knockout of this gene causes progressive hind limb paralysis in mice ((46,47). The enigmatic *Dclk1* (doublecortin-like kinase 1) implicated in growth dysregulation in several cancers (48) is part of this cluster and public array data in mouse (biogps.org) reveal the transcript is greatly over-expressed in C2C12 myoblasts. The cluster also contains known regulatory growth factors *Fgf7, Tgfb2* and downstream target *Fst*. Finally, the cluster contains transcripts encoding enzymes of polyamine synthesis (*Odc, Sms*), which regulates cell proliferation in myogenesis (49)

Cluster 7 contains transcripts encoding multiple muscle-expressed intermediate filament (Krt) proteins (but not desmin), junction-associated proteins and cell adhesion molecules with known function in skeletal muscle integrity and force transductions including several desmoglein (Dsg) and desmocollin (Dsc) genes and desmoplakin (Dsp) that combine to form desmosomes (50). The clear separation of this cluster indicates that structural integrity of skeletal muscle is independently regulated. Interestingly, Cluster 7 contains all three members of the grainyhead-like family (*Grhl1/2/3*) which also regulate expression of junction-associated transcripts in epithelia (51)

There are three separate connective tissue clusters associated with distinct collagen subunits, each with associated specific transcription factors. The smallest includes *Fbn1*, the gene associated with Marfan syndrome, which has a specific function in elastic fibres (52) as well as multiple members of the Adamts family (53)

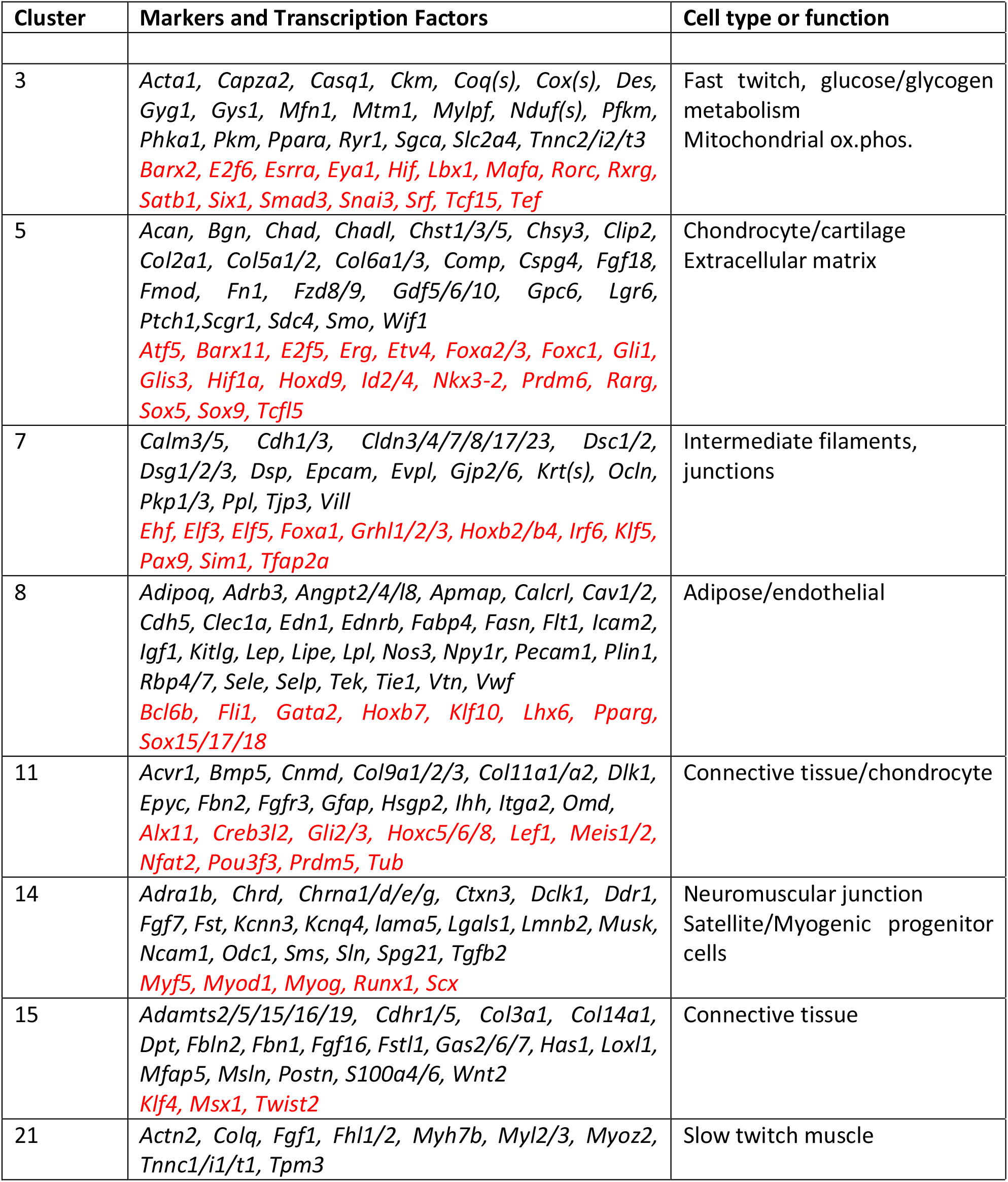

## References

1. Smith, J.R., Hayman, G.T., Wang, S.J., Laulederkind, S.J.F., Hoffman, M.J., Kaldunski, M.L., Tutaj, M., Thota, J., Nalabolu, H.S., Ellanki, S.L.R. et al. (2020) The Year of the Rat: The Rat Genome Database at 20: a multi-species knowledgebase and analysis platform. Nucleic Acids Res, 48, D731–D742.

2. Gibbs, R.A., Weinstock, G.M., Metzker, M.L., Muzny, D.M., Sodergren, E.J., Scherer, S., Scott, G., Steffen, D., Worley, K.C., Burch, P.E. et al. (2004) Genome sequence of the Brown Norway rat yields insights into mammalian evolution. Nature, 428, 493–521.

3. Atanur, S.S., Diaz, A.G., Maratou, K., Sarkis, A., Rotival, M., Game, L., Tschannen, M.R., Kaisaki, P.J., Otto, G.W., Ma, M.C. et al. (2013) Genome sequencing reveals loci under artificial selection that underlie disease phenotypes in the laboratory rat. Cell, 154, 691–703.

4. Szpirer, C. (2020) Rat models of human diseases and related phenotypes: a systematic inventory of the causative genes. J Biomed Sci, 27, 84.

5. Consortium, G.T. (2015) Human genomics. The Genotype-Tissue Expression (GTEx) pilot analysis: multitissue gene regulation in humans. Science, 348, 648–660.

6. Consortium, F., the, R.P., Clst Forrest, A.R., Kawaji, H., Rehli, M., Baillie, J.K., de Hoon, M.J., Haberle, V., Lassmann, T. et al. (2014) A promoter-level mammalian expression atlas. Nature, 507, 462–470.

7. Bush, S.J., Freem, L., MacCallum, A.J., O’Dell, J., Wu, C., Afrasiabi, C., Psifidi, A., Stevens, M.P., Smith, J., Summers, K.M. et al. (2018) Combination of novel and public RNA-seq datasets to generate an mRNA expression atlas for the domestic chicken. BMC Genomics, 19, 594.

8. Clark, E.L., Bush, S.J., McCulloch, M.E.B., Farquhar, I.L., Young, R., Lefevre, L., Pridans, C., Tsang, H.G., Wu, C., Afrasiabi, C. et al. (2017) A high resolution atlas of gene expression in the domestic sheep (Ovis aries). PLoS Genet, 13, e1006997.

9. Young, R., Lefevre, L., Bush, S.J., Joshi, A., Singh, S.H., Jadhav, S.K., Dhanikachalam, V., Lisowski, Z.M., Iamartino, D., Summers, K.M. et al. (2019) A Gene Expression Atlas of the Domestic Water Buffalo (Bubalus bubalis). Front Genet, 10, 668.

10. Muriuki, C., Bush, S.J., Salavati, M., McCulloch, M.E.B., Lisowski, Z.M., Agaba, M., Djikeng, A., Hume, D.A. and Clark, E.L. (2019) A Mini-Atlas of Gene Expression for the Domestic Goat (Capra hircus). Front Genet, 10, 1080.

11. Summers, K.M., Bush, S.J., Wu, C., Su, A.I., Muriuki, C., Clark, E.L., Finlayson, H.A., Eory, L., Waddell, L.A., Talbot, R. et al. (2019) Functional Annotation of the Transcriptome of the Pig, Sus scrofa, Based Upon Network Analysis of an RNAseq Transcriptional Atlas. Front Genet, 10, 1355.

12. Gillis, J. and Pavlidis, P. (2012) “Guilt by association” is the exception rather than the rule in gene networks. PLoS Comput Biol, 8, e1002444.

13. Ballouz, S., Weber, M., Pavlidis, P. and Gillis, J. (2017) EGAD: ultra-fast functional analysis of gene networks. Bioinformatics, 33, 612–614.

14. Freeman, T.C., Ivens, A., Baillie, J.K., Beraldi, D., Barnett, M.W., Dorward, D., Downing, A., Fairbairn, L., Kapetanovic, R., Raza, S. et al. (2012) A gene expression atlas of the domestic pig. BMC Biol, 10, 90.

15. Giotti, B., Chen, S.H., Barnett, M.W., Regan, T., Ly, T., Wiemann, S., Hume, D.A. and Freeman, T.C. (2018) Assembly of a Parts List of the Human Mitotic Cell Cycle Machinery. J Mol Cell Biol.

16. Hume, D.A., Summers, K.M., Raza, S., Baillie, J.K. and Freeman, T.C. (2010) Functional clustering and lineage markers: insights into cellular differentiation and gene function from large-scale microarray studies of purified primary cell populations. Genomics, 95, 328–338.

17. Mabbott, N.A., Baillie, J.K., Brown, H., Freeman, T.C. and Hume, D.A. (2013) An expression atlas of human primary cells: inference of gene function from coexpression networks. BMC Genomics, 14, 632.

18. Singh, A.J., Ramsey, S.A., Filtz, T.M. and Kioussi, C. (2018) Differential gene regulatory networks in development and disease. Cell Mol Life Sci, 75, 1013–1025.

19. Doig, T.N., Hume, D.A., Theocharidis, T., Goodlad, J.R., Gregory, C.D. and Freeman, T.C. (2013) Coexpression analysis of large cancer datasets provides insight into the cellular phenotypes of the tumour microenvironment. BMC Genomics, 14, 469.

20. Summers, K.M., Bush, S.J. and Hume, D.A. (2020) Network analysis of transcriptomic diversity amongst resident tissue macrophages and dendritic cells in the mouse mononuclear phagocyte system. PLoS Biol, 18, e3000859.

21. Jubb, A.W., Young, R.S., Hume, D.A. and Bickmore, W.A. (2016) Enhancer Turnover Is Associated with a Divergent Transcriptional Response to Glucocorticoid in Mouse and Human Macrophages. J Immunol, 196, 813–822.

22. Villar, D., Berthelot, C., Aldridge, S., Rayner, T.F., Lukk, M., Pignatelli, M., Park, T.J., Deaville, R., Erichsen, J.T., Jasinska, A.J. et al. (2015) Enhancer evolution across 20 mammalian species. Cell, 160, 554–566.

23. Ji, X., Li, P., Fuscoe, J.C., Chen, G., Xiao, W., Shi, L., Ning, B., Liu, Z., Hong, H., Wu, J. et al. (2020) A comprehensive rat transcriptome built from large scale RNA-seq-based annotation. Nucleic Acids Res, 48, 8320–8331.

24. Sollner, J.F., Leparc, G., Hildebrandt, T., Klein, H., Thomas, L., Stupka, E. and Simon, E. (2017) An RNA-Seq atlas of gene expression in mouse and rat normal tissues. Sci Data, 4, 170185.

25. Wang, X., You, X., Langer, J.D., Hou, J., Rupprecht, F., Vlatkovic, I., Quedenau, C., Tushev, G., Epstein, I., Schaefke, B. et al. (2019) Full-length transcriptome reconstruction reveals a large diversity of RNA and protein isoforms in rat hippocampus. Nat Commun, 10, 5009.

26. Bray, N.L., Pimentel, H., Melsted, P. and Pachter, L. (2016) Near-optimal probabilistic RNA-seq quantification. Nat Biotechnol, 34, 525–527.

27. Choudhary, S. (2019) pysradb: A Python package to query next-generation sequencing metadata and data from NCBI Sequence Read Archive. F1000Res, 8, 532.

28. Wu, C., Jin, X., Tsueng, G., Afrasiabi, C. and Su, A.I. (2016) BioGPS: building your own mash-up of gene annotations and expression profiles. Nucleic Acids Res, 44, D313–316.

29. Wu, C., Orozco, C., Boyer, J., Leglise, M., Goodale, J., Batalov, S., Hodge, C.L., Haase, J., Janes, J., Huss, J.W., 3rd et al. (2009) BioGPS: an extensible and customizable portal for querying and organizing gene annotation resources. Genome Biol, 10, R130.

30. Theocharidis, A., van Dongen, S., Enright, A.J. and Freeman, T.C. (2009) Network visualization and analysis of gene expression data using BioLayout Express(3D). Nat Protoc, 4, 1535–1550.

31. van Dongen, S. and Abreu-Goodger, C. (2012) In Van Helden, J., Toussaint, A. and Theiffry, D. (eds.), Bacterial molecular networks: Mtehods and protocols. Springer, New York, NY, USA.

32. Yu, Y., Fuscoe, J.C., Zhao, C., Guo, C., Jia, M., Qing, T., Bannon, D.I., Lancashire, L., Bao, W., Du, T. et al. (2014) A rat RNA-Seq transcriptomic BodyMap across 11 organs and 4 developmental stages. Nat Commun, 5, 3230.

33. Grimes, S.R. (2004) Testis-specific transcriptional control. Gene, 343, 11–22.

34. Okitsu, Y., Nagano, M., Yamagata, T., Ito, C., Toshimori, K., Dohra, H., Fujii, W. and Yogo, K. (2020) Dlec1 is required for spermatogenesis and male fertility in mice. Sci Rep, 10, 18883.

35. Li, M., Zheng, J., Li, G., Lin, Z., Li, D., Liu, D., Feng, H., Cao, D., Ng, E.H.Y., Li, R.H.W. et al. (2021) The male germline-specific protein MAPS is indispensable for pachynema progression and fertility. Proc Natl Acad Sci U S A, 118.

36. Wu, X.L., Yun, D.M., Gao, S., Liang, A.J., Duan, Z.Z., Wang, H.S., Wang, G.S. and Sun, F. (2020) The testis-specific gene 1700102P08Rik is essential for male fertility. Mol Reprod Dev, 87, 231–240.

37. Cunningham, D.B., Segretain, D., Arnaud, D., Rogner, U.C. and Avner, P. (1998) The mouse Tsx gene is expressed in Sertoli cells of the adult testis and transiently in premeiotic germ cells during puberty. Dev Biol, 204, 345–360.

38. Svingen, T., Beverdam, A., Verma, P., Wilhelm, D. and Koopman, P. (2007) Aard is specifically up-regulated in Sertoli cells during mouse testis differentiation. Int J Dev Biol, 51, 255–258.

39. Akinloye, O., Gromoll, J., Callies, C., Nieschlag, E. and Simoni, M. (2007) Mutation analysis of the X-chromosome linked, testis-specific TAF7L gene in spermatogenic failure. Andrologia, 39, 190–195.

40. Cheng, Y., Buffone, M.G., Kouadio, M., Goodheart, M., Page, D.C., Gerton, G.L., Davidson, I. and Wang, P.J. (2007) Abnormal sperm in mice lacking the Taf7l gene. Mol Cell Biol, 27, 2582–2589.

41. Chang, E., Fu, C., Coon, S.L., Alon, S., Bozinoski, M., Breymaier, M., Bustos, D.M., Clokie, S.J., Gothilf, Y., Esnault, C. et al. (2020) Resource: A multi-species multi-timepoint transcriptome database and webpage for the pineal gland and retina. J Pineal Res, 69, e12673.

42. Rohde, K., Rovsing, L., Ho, A.K., Moller, M. and Rath, M.F. (2014) Circadian dynamics of the cone-rod homeobox (CRX) transcription factor in the rat pineal gland and its role in regulation of arylalkylamine N-acetyltransferase (AANAT). Endocrinology, 155, 2966–2975.

43. Bailey, M.J., Coon, S.L., Carter, D.A., Humphries, A., Kim, J.S., Shi, Q., Gaildrat, P., Morin, F., Ganguly, S., Hogenesch, J.B. et al. (2009) Night/day changes in pineal expression of >600 genes: central role of adrenergic/cAMP signaling. J Biol Chem, 284, 7606–7622.

44. Goding, C.R. and Arnheiter, H. (2019) MITF-the first 25 years. Genes Dev, 33, 983–1007.

45. Fraser, A.M. and Davey, M.G. (2019) TALPID3 in Joubert syndrome and related ciliopathy disorders. Curr Opin Genet Dev, 56, 41–48.

46. Morishita, H. and Yagi, T. (2007) Protocadherin family: diversity, structure, and function. Curr Opin Cell Biol, 19, 584–592.

47. Missaglia, S., Tavian, D. and Angelini, C. (2021) ETF dehydrogenase advances in molecular genetics and impact on treatment. Crit Rev Biochem Mol Biol, 56, 360–372.

48. Chen, C.M., Wang, H.Y., You, L.R., Shang, R.L. and Liu, F.C. (2010) Expression analysis of an evolutionarily conserved metallophosphodiesterase gene, Mpped1, in the normal and beta-catenin-deficient malformed dorsal telencephalon. Dev Dyn, 239, 1797–1806.

49. Maubert, E., Slama, A., Ciofi, P., Viollet, C., Tramu, G., Dupouy, J.P. and Epelbaum, J. (1994) Developmental patterns of somatostatin-receptors and somatostatin-immunoreactivity during early neurogenesis in the rat. Neuroscience, 62, 317–325.

50. Chu, C.H., Chen, J.S., Chuang, P.C., Su, C.H., Chan, Y.L., Yang, Y.J., Chiang, Y.T., Su, Y.Y., Gean, P.W. and Sun, H.S. (2020) TIAM2S as a novel regulator for serotonin level enhances brain plasticity and locomotion behavior. FASEB J, 34, 3267–3288.

51. Stolt, C.C., Lommes, P., Friedrich, R.P. and Wegner, M. (2004) Transcription factors Sox8 and Sox10 perform non-equivalent roles during oligodendrocyte development despite functional redundancy. Development, 131, 2349–2358.

52. Artegiani, B., Lyubimova, A., Muraro, M., van Es, J.H., van Oudenaarden, A. and Clevers, H. (2017) A Single-Cell RNA Sequencing Study Reveals Cellular and Molecular Dynamics of the Hippocampal Neurogenic Niche. Cell Rep, 21, 3271–3284.

53. Perez-Frances, M., van Gurp, L., Abate, M.V., Cigliola, V., Furuyama, K., Bru-Tari, E., Oropeza, D., Carreaux, T., Fujitani, Y., Thorel, F. et al. (2021) Pancreatic Ppy-expressing gamma-cells display mixed phenotypic traits and the adaptive plasticity to engage insulin production. Nat Commun, 12, 4458.

54. Davis, M.R., Arner, E., Duffy, C.R., De Sousa, P.A., Dahlman, I., Arner, P. and Summers, K.M. (2016) Expression of FBN1 during adipogenesis: Relevance to the lipodystrophy phenotype in Marfan syndrome and related conditions. Mol Genet Metab, 119, 174–185.

55. Attanasio, M., Pratelli, E., Porciani, M.C., Evangelisti, L., Torricelli, E., Pellicano, G., Abbate, R., Gensini, G.F. and Pepe, G. (2013) Dural ectasia and FBN1 mutation screening of 40 patients with Marfan syndrome and related disorders: role of dural ectasia for the diagnosis. Eur J Med Genet, 56, 356–360.

56. Jespersen, K., Liu, Z., Li, C., Harding, P., Sestak, K., Batra, R., Stephenson, C.A., Foley, R.T., Greene, H., Meisinger, T. et al. (2020) Enhanced Notch3 signaling contributes to pulmonary emphysema in a Murine Model of Marfan syndrome. Sci Rep, 10, 10949.

57. Anderson, J.L., Head, S.I., Rae, C. and Morley, J.W. (2002) Brain function in Duchenne muscular dystrophy. Brain, 125, 4–13.

58. O’Rourke, J.G., Bogdanik, L., Yanez, A., Lall, D., Wolf, A.J., Muhammad, A.K., Ho, R., Carmona, S., Vit, J.P., Zarrow, J. et al. (2016) C9orf72 is required for proper macrophage and microglial function in mice. Science, 351, 1324–1329.

59. Walsh, N.C., Cahill, M., Carninci, P., Kawai, J., Okazaki, Y., Hayashizaki, Y., Hume, D.A. and Cassady, A.I. (2003) Multiple tissue-specific promoters control expression of the murine tartrate-resistant acid phosphatase gene. Gene, 307, 111–123.

60. Mitic, N., Valizadeh, M., Leung, E.W., de Jersey, J., Hamilton, S., Hume, D.A., Cassady, A.I. and Schenk, G. (2005) Human tartrate-resistant acid phosphatase becomes an effective ATPase upon proteolytic activation. Arch Biochem Biophys, 439, 154–164.

61. Lang, P., van Harmelen, V., Ryden, M., Kaaman, M., Parini, P., Carneheim, C., Cassady, A.I., Hume, D.A., Andersson, G. and Arner, P. (2008) Monomeric tartrate resistant acid phosphatase induces insulin sensitive obesity. PLoS One, 3, e1713.

62. Sengupta, A., Rhoades, S.D., Kim, E.J., Nayak, S., Grant, G.R., Meerlo, P. and Weljie, A.M. (2017) Sleep restriction induced energy, methylation and lipogenesis metabolic switches in rat liver. Int J Biochem Cell Biol, 93, 129–135.

63. Huang, X., Cai, H., Ammar, R., Zhang, Y., Wang, Y., Ravi, K., Thompson, J. and Jarai, G. (2019) Molecular characterization of a precision-cut rat liver slice model for the evaluation of antifibrotic compounds. Am J Physiol Gastrointest Liver Physiol, 316, G15–G24.

64. Kimball, S.R., Horetsky, R.L. and Jefferson, L.S. (1995) Hormonal regulation of albumin gene expression in primary cultures of rat hepatocytes. Am J Physiol, 268, E6–14.

65. Qvartskhava, N., Lang, P.A., Gorg, B., Pozdeev, V.I., Ortiz, M.P., Lang, K.S., Bidmon, H.J., Lang, E., Leibrock, C.B., Herebian, D. et al. (2015) Hyperammonemia in gene-targeted mice lacking functional hepatic glutamine synthetase. Proc Natl Acad Sci U S A, 112, 5521–5526.

66. Pridans, C., Irvine, K.M., Davis, G.M., Lefevre, L., Bush, S.J. and Hume, D.A. (2020) Transcriptomic Analysis of Rat Macrophages. Front Immunol, 11, 594594.

67. Hume, D.A., Caruso, M., Keshvari, S., Patkar, O.L., Sehgal, A., Bush, S.J., Summers, K.M., Pridans, C. and Irvine, K.M. (2021) The Mononuclear Phagocyte System of the Rat. J Immunol, 206, 2251–2263.

68. Patkar, O.L., Caruso, M., Teakle, N., Keshvari, S., Bush, S.J., Pridans, C., Belmer, A., Summers, K.M., Irvine, K.M. and Hume, D.A. (2021) Analysis of homozygous and heterozygous Csf1r knockout in the rat as a model for understanding microglial function in brain development and the impacts of human CSF1R mutations. Neurobiol Dis, 151, 105268.

69. Kohyama, M., Ise, W., Edelson, B.T., Wilker, P.R., Hildner, K., Mejia, C., Frazier, W.A., Murphy, T.L. and Murphy, K.M. (2009) Role for Spi-C in the development of red pulp macrophages and splenic iron homeostasis. Nature, 457, 318–321.

70. Rojo, R., Pridans, C., Langlais, D. and Hume, D.A. (2017) Transcriptional mechanisms that control expression of the macrophage colony-stimulating factor receptor locus. Clin Sci (Lond), 131, 2161–2182.

71. Irvine, K.M., Caruso, M., Cestari, M.F., Davis, G.M., Keshvari, S., Sehgal, A., Pridans, C. and Hume, D.A. (2020) Analysis of the impact of CSF-1 administration in adult rats using a novel Csf1r-mApple reporter gene. J Leukoc Biol, 107, 221–235.

72. Summers, K.M. and Hume, D.A. (2017) Identification of the macrophage-specific promoter signature in FANTOM5 mouse embryo developmental time course data. J Leukoc Biol, 102, 1081–1092.

73. Ben-Moshe, S. and Itzkovitz, S. (2019) Spatial heterogeneity in the mammalian liver. Nat Rev Gastroenterol Hepatol, 16, 395–410.

74. Halpern, K.B., Shenhav, R., Matcovitch-Natan, O., Toth, B., Lemze, D., Golan, M., Massasa, E.E., Baydatch, S., Landen, S., Moor, A.E. et al. (2017) Single-cell spatial reconstruction reveals global division of labour in the mammalian liver. Nature, 542, 352–356.

75. Atger, F., Gobet, C., Marquis, J., Martin, E., Wang, J., Weger, B., Lefebvre, G., Descombes, P., Naef, F. and Gachon, F. (2015) Circadian and feeding rhythms differentially affect rhythmic mRNA transcription and translation in mouse liver. Proc Natl Acad Sci U S A, 112, E6579–6588.

76. Cheng, X., Kim, S.Y., Okamoto, H., Xin, Y., Yancopoulos, G.D., Murphy, A.J. and Gromada, J. (2018) Glucagon contributes to liver zonation. Proc Natl Acad Sci U S A, 115, E4111–E4119.

77. Heidenreich, S., Weber, P., Stephanowitz, H., Petricek, K.M., Schutte, T., Oster, M., Salo, A.M., Knauer, M., Goehring, I., Yang, N. et al. (2020) The glucose-sensing transcription factor ChREBP is targeted by proline hydroxylation. J Biol Chem, 295, 17158–17168.

78. Jiang, Y., Feng, D., Ma, X., Fan, S., Gao, Y., Fu, K., Wang, Y., Sun, J., Yao, X., Liu, C. et al. (2019) Pregnane X Receptor Regulates Liver Size and Liver Cell Fate by Yes-Associated Protein Activation in Mice. Hepatology, 69, 343–358.

79. Gialitakis, M., Tolaini, M., Li, Y., Pardo, M., Yu, L., Toribio, A., Choudhary, J.S., Niakan, K., Papayannopoulos, V. and Stockinger, B. (2017) Activation of the Aryl Hydrocarbon Receptor Interferes with Early Embryonic Development. Stem Cell Reports, 9, 1377–1386.

80. Taniguchi, M. and Yoshida, H. (2017) TFE3, HSP47, and CREB3 Pathways of the Mammalian Golgi Stress Response. Cell Struct Funct, 42, 27–36.

81. Gosselin, D., Skola, D., Coufal, N.G., Holtman, I.R., Schlachetzki, J.C.M., Sajti, E., Jaeger, B.N., O’Connor, C., Fitzpatrick, C., Pasillas, M.P. et al. (2017) An environment-dependent transcriptional network specifies human microglia identity. Science, 356.

82. Keshvari, S., Caruso, M., Teakle, N., Batoon, L., Sehgal, A., Patkar, O.L., Ferrari-Cestari, M., Snell, C.E., Chen, C., Stevenson, A. et al. (2021) CSF1R-dependent macrophages control postnatal somatic growth and organ maturation. PLoS Genet, 17, e1009605.

83. Niederkorn, M., Agarwal, P. and Starczynowski, D.T. (2020) TIFA and TIFAB: FHA-domain proteins involved in inflammation, hematopoiesis, and disease. Exp Hematol, 90, 18–29.

84. Conforto, T.L., Zhang, Y., Sherman, J. and Waxman, D.J. (2012) Impact of CUX2 on the female mouse liver transcriptome: activation of female-biased genes and repression of male-biased genes. Mol Cell Biol, 32, 4611–4627.

85. Lau-Corona, D., Suvorov, A. and Waxman, D.J. (2017) Feminization of Male Mouse Liver by Persistent Growth Hormone Stimulation: Activation of Sex-Biased Transcriptional Networks and Dynamic Changes in Chromatin States. Mol Cell Biol, 37.

86. Lahnemann, D., Koster, J., Szczurek, E., McCarthy, D.J., Hicks, S.C., Robinson, M.D., Vallejos, C.A., Campbell, K.R., Beerenwinkel, N., Mahfouz, A. et al. (2020) Eleven grand challenges in single-cell data science. Genome Biol, 21, 31.

## References

1. Patkar, O.L., Caruso, M., Teakle, N., Keshvari, S., Bush, S.J., Pridans, C., Belmer, A., Summers, K.M., Irvine, K.M. and Hume, D.A. (2021) Analysis of homozygous and heterozygous Csf1r knockout in the rat as a model for understanding microglial function in brain development and the impacts of human CSF1R mutations. Neurobiol Dis, 151, 105268.

2. Faure, L., Wang, Y., Kastriti, M.E., Fontanet, P., Cheung, K.K.Y., Petitpre, C., Wu, H., Sun, L.L., Runge, K., Croci, L. et al. (2020) Single cell RNA sequencing identifies early diversity of sensory neurons forming via bi-potential intermediates. Nat Commun, 11, 4175.

3. Santiago, C. and Bashaw, G.J. (2014) Transcription factors and effectors that regulate neuronal morphology. Development, 141, 4667–4680.

4. Neumann, B., Baror, R., Zhao, C., Segel, M., Dietmann, S., Rawji, K.S., Foerster, S., McClain, C.R., Chalut, K., van Wijngaarden, P. et al. (2019) Metformin Restores CNS Remyelination Capacity by Rejuvenating Aged Stem Cells. Cell Stem Cell, 25, 473–485 e478.

5. Ji, Y., Shen, M., Wang, X., Zhang, S., Yu, S., Chen, G., Gu, X. and Ding, F. (2012) Comparative proteomic analysis of primary schwann cells and a spontaneously immortalized schwann cell line RSC 96: a comprehensive overview with a focus on cell adhesion and migration related proteins. J Proteome Res, 11, 3186–3198.

6. Park, S.K., Miller, R., Krane, I. and Vartanian, T. (2001) The erbB2 gene is required for the development of terminally differentiated spinal cord oligodendrocytes. J Cell Biol, 154, 1245–1258.

7. Jessen, K.R., Mirsky, R. and Lloyd, A.C. (2015) Schwann Cells: Development and Role in Nerve Repair. Cold Spring Harb Perspect Biol, 7, a020487.

8. Mays, J.C., Kelly, M.C., Coon, S.L., Holtzclaw, L., Rath, M.F., Kelley, M.W. and Klein, D.C. (2018) Single-cell RNA sequencing of the mammalian pineal gland identifies two pinealocyte subtypes and cell type-specific daily patterns of gene expression. PLoS One, 13, e0205883.

9. Hasel, P., Rose, I.V.L., Sadick, J.S., Kim, R.D. and Liddelow, S.A. (2021) Neuroinflammatory astrocyte subtypes in the mouse brain. Nat Neurosci.

10. Consortium, F., the, R.P., Clst Forrest, A.R., Kawaji, H., Rehli, M., Baillie, J.K., de Hoon, M.J., Haberle, V., Lassmann, T. et al. (2014) A promoter-level mammalian expression atlas. Nature, 507, 462–470.

11. Krupenko, N.I., Sharma, J., Fogle, H.M., Pediaditakis, P., Strickland, K.C., Du, X., Helke, K.L., Sumner, S. and Krupenko, S.A. (2021) Knockout of Putative Tumor Suppressor Aldh1l1 in Mice Reprograms Metabolism to Accelerate Growth of Tumors in a Diethylnitrosamine (DEN) Model of Liver Carcinogenesis. Cancers (Basel), 13.

12. Batiuk, M.Y., Martirosyan, A., Wahis, J., de Vin, F., Marneffe, C., Kusserow, C., Koeppen, J., Viana, J.F., Oliveira, J.F., Voet, T. et al. (2020) Identification of region-specific astrocyte subtypes at single cell resolution. Nat Commun, 11, 1220.

13. Walz, W. and Lang, M.K. (1998) Immunocytochemical evidence for a distinct GFAP-negative subpopulation of astrocytes in the adult rat hippocampus. Neurosci Lett, 257, 127–130.

14. Nouel, D., Sarret, P., Vincent, J.P., Mazella, J. and Beaudet, A. (1999) Pharmacological, molecular and functional characterization of glial neurotensin receptors. Neuroscience, 94, 1189–1197.

15. Novielli-Kuntz, N.M., Press, E.R., Barr, K., Prado, M.A.M. and Laird, D.W. (2021) Mutant Cx30-A88V mice exhibit hydrocephaly and sex-dependent behavioral abnormalities, implicating a functional role for Cx30 in the brain. Dis Model Mech, 14.

16. Sandoval, M., Luarte, A., Herrera-Molina, R., Varas-Godoy, M., Santibanez, M., Rubio, F.J., Smit, A.B., Gundelfinger, E.D., Li, K.W., Smalla, K.H. et al. (2013) The glycolytic enzyme aldolase C is up-regulated in rat forebrain microsomes and in the cerebrospinal fluid after repetitive fluoxetine treatment. Brain Res, 1520, 1–14.

17. Fry, A.E., Cushion, T.D. and Pilz, D.T. (2014) The genetics of lissencephaly. Am J Med Genet C Semin Med Genet, 166C, 198–210.

18. Lee, J.W., Chou, C.L. and Knepper, M.A. (2015) Deep Sequencing in Microdissected Renal Tubules Identifies Nephron Segment-Specific Transcriptomes. J Am Soc Nephrol, 26, 2669–2677.

19. Clark, J.Z., Chen, L., Chou, C.L., Jung, H.J., Lee, J.W. and Knepper, M.A. (2019) Representation and relative abundance of cell-type selective markers in whole-kidney RNA-Seq data. Kidney Int, 95, 787–796.

20. Wang, X., Park, J., Susztak, K., Zhang, N.R. and Li, M. (2019) Bulk tissue cell type deconvolution with multi-subject single-cell expression reference. Nat Commun, 10, 380.

21. Sheng, X., Guan, Y., Ma, Z., Wu, J., Liu, H., Qiu, C., Vitale, S., Miao, Z., Seasock, M.J., Palmer, M. et al. (2021) Mapping the genetic architecture of human traits to cell types in the kidney identifies mechanisms of disease and potential treatments. Nat Genet, 53, 1322–1333.

22. Nakajima, K., Miranda, A., Craig, D.W., Shekhtman, T., Kmoch, S., Bleyer, A., Szelinger, S., Kato, T. and Kelsoe, J.R. (2020) Ntrk1 mutation co-segregating with bipolar disorder and inherited kidney disease in a multiplex family causes defects in neuronal growth and depression-like behavior in mice. Transl Psychiatry, 10, 407.

23. Chen, L., Lee, J.W., Chou, C.L., Nair, A.V., Battistone, M.A., Paunescu, T.G., Merkulova, M., Breton, S., Verlander, J.W., Wall, S.M. et al. (2017) Transcriptomes of major renal collecting duct cell types in mouse identified by single-cell RNA-seq. Proc Natl Acad Sci U S A, 114, E9989–E9998.

24. Grabert, K., Sehgal, A., Irvine, K.M., Wollscheid-Lengeling, E., Ozdemir, D.D., Stables, J., Luke, G.A., Ryan, M.D., Adamson, A., Humphreys, N.E. et al. (2020) A Transgenic Line That Reports CSF1R Protein Expression Provides a Definitive Marker for the Mouse Mononuclear Phagocyte System. J Immunol, 205, 3154–3166.

25. Irvine, K.M., Caruso, M., Cestari, M.F., Davis, G.M., Keshvari, S., Sehgal, A., Pridans, C. and Hume, D.A. (2020) Analysis of the impact of CSF-1 administration in adult rats using a novel Csf1r-mApple reporter gene. J Leukoc Biol, 107, 221–235.

26. Hume, D.A. and Gordon, S. (1983) Mononuclear phagocyte system of the mouse defined by immunohistochemical localization of antigen F4/80. Identification of resident macrophages in renal medullary and cortical interstitium and the juxtaglomerular complex. J Exp Med, 157, 1704–1709.

27. Zimmerman, K.A., Yang, Z., Lever, J.M., Li, Z., Croyle, M.J., Agarwal, A., Yoder, B.K. and George, J.F. (2021) Kidney resident macrophages in the rat have minimal turnover and replacement by blood monocytes. Am J Physiol Renal Physiol, 321, F162–F169.

28. Paik, D.T., Cho, S., Tian, L., Chang, H.Y. and Wu, J.C. (2020) Single-cell RNA sequencing in cardiovascular development, disease and medicine. Nat Rev Cardiol, 17, 457–473.

29. Farbehi, N., Patrick, R., Dorison, A., Xaymardan, M., Janbandhu, V., Wystub-Lis, K., Ho, J.W., Nordon, R.E. and Harvey, R.P. (2019) Single-cell expression profiling reveals dynamic flux of cardiac stromal, vascular and immune cells in health and injury. Elife, 8.

30. Golan-Lagziel, T., Lewis, Y.E., Shkedi, O., Douvdevany, G., Caspi, L.H. and Kehat, I. (2018) Analysis of rat cardiac myocytes and fibroblasts identifies combinatorial enhancer organization and transcription factor families. J Mol Cell Cardiol, 116, 91–105.

31. Mahmoud, A.I., Kocabas, F., Muralidhar, S.A., Kimura, W., Koura, A.S., Thet, S., Porrello, E.R. and Sadek, H.A. (2013) Meis1 regulates postnatal cardiomyocyte cell cycle arrest. Nature, 497, 249–253.

32. Parvatiyar, M.S., Brownstein, A.J., Kanashiro-Takeuchi, R.M., Collado, J.R., Dieseldorff Jones, K.M., Gopal, J., Hammond, K.G., Marshall, J.L., Ferrel, A., Beedle, A.M. et al. (2019) Stabilization of the cardiac sarcolemma by sarcospan rescues DMD-associated cardiomyopathy. JCI Insight, 5.

33. Coral-Vazquez, R., Cohn, R.D., Moore, S.A., Hill, J.A., Weiss, R.M., Davisson, R.L., Straub, V., Barresi, R., Bansal, D., Hrstka, R.F. et al. (1999) Disruption of the sarcoglycan-sarcospan complex in vascular smooth muscle: a novel mechanism for cardiomyopathy and muscular dystrophy. Cell, 98, 465–474.

34. Weinberger, T., Esfandyari, D., Messerer, D., Percin, G., Schleifer, C., Thaler, R., Liu, L., Stremmel, C., Schneider, V., Vagnozzi, R.J. et al. (2020) Ontogeny of arterial macrophages defines their functions in homeostasis and inflammation. Nat Commun, 11, 4549.

35. Rodero, M.P., Poupel, L., Loyher, P.L., Hamon, P., Licata, F., Pessel, C., Hume, D.A., Combadiere, C. and Boissonnas, A. (2015) Immune surveillance of the lung by migrating tissue monocytes. Elife, 4, e07847.

36. Hume, D.A. and Freeman, T.C. (2014) Transcriptomic analysis of mononuclear phagocyte differentiation and activation. Immunol Rev, 262, 74–84.

37. Fedele, L. and Brand, T. (2020) The Intrinsic Cardiac Nervous System and Its Role in Cardiac Pacemaking and Conduction. J Cardiovasc Dev Dis, 7.

38. Werner, P., Paluru, P., Simpson, A.M., Latney, B., Iyer, R., Brodeur, G.M. and Goldmuntz, E. (2014) Mutations in NTRK3 suggest a novel signaling pathway in human congenital heart disease. Hum Mutat, 35, 1459–1468.

39. Van Dyke, J.M., Smit-Oistad, I.M., Macrander, C., Krakora, D., Meyer, M.G. and Suzuki, M. (2016) Macrophage-mediated inflammation and glial response in the skeletal muscle of a rat model of familial amyotrophic lateral sclerosis (ALS). Exp Neurol, 277, 275–282.

40. Liu, W., Klose, A., Forman, S., Paris, N.D., Wei-LaPierre, L., Cortes-Lopez, M., Tan, A., Flaherty, M., Miura, P., Dirksen, R.T. et al. (2017) Loss of adult skeletal muscle stem cells drives age-related neuromuscular junction degeneration. Elife, 6.

41. Rodriguez Cruz, P.M., Cossins, J., Beeson, D. and Vincent, A. (2020) The Neuromuscular Junction in Health and Disease: Molecular Mechanisms Governing Synaptic Formation and Homeostasis. Front Mol Neurosci, 13, 610964.

42. Umansky, K.B., Gruenbaum-Cohen, Y., Tsoory, M., Feldmesser, E., Goldenberg, D., Brenner, O. and Groner, Y. (2015) Runx1 Transcription Factor Is Required for Myoblasts Proliferation during Muscle Regeneration. PLoS Genet, 11, e1005457.

43. Blitz, E., Sharir, A., Akiyama, H. and Zelzer, E. (2013) Tendon-bone attachment unit is formed modularly by a distinct pool of Scx-and Sox9-positive progenitors. Development, 140, 2680–2690.

44. Seale, P., Sabourin, L.A., Girgis-Gabardo, A., Mansouri, A., Gruss, P. and Rudnicki, M.A. (2000) Pax7 is required for the specification of myogenic satellite cells. Cell, 102, 777–786.

45. Rossi, C.A., Pozzobon, M., Ditadi, A., Archacka, K., Gastaldello, A., Sanna, M., Franzin, C., Malerba, A., Milan, G., Cananzi, M. et al. (2010) Clonal characterization of rat muscle satellite cells: proliferation, metabolism and differentiation define an intrinsic heterogeneity. PLoS One, 5, e8523.

46. Davenport, A., Bivona, A., Latson, W., Lemanski, L.F. and Cheriyath, V. (2016) Loss of Maspardin Attenuates the Growth and Maturation of Mouse Cortical Neurons. Neurodegener Dis, 16, 260–272.

47. Soderblom, C., Stadler, J., Jupille, H., Blackstone, C., Shupliakov, O. and Hanna, M.C. (2010) Targeted disruption of the Mast syndrome gene SPG21 in mice impairs hind limb function and alters axon branching in cultured cortical neurons. Neurogenetics, 11, 369–378.

48. Agulto, R.L., Rogers, M.M., Tan, T.C., Ramkumar, A., Downing, A.M., Bodin, H., Castro, J., Nowakowski, D.W. and Ori-McKenney, K.M. (2021) Autoregulatory control of microtubule binding in doublecortin-like kinase 1. Elife, 10.

49. Machado, L., Geara, P., Camps, J., Dos Santos, M., Teixeira-Clerc, F., Van Herck, J., Varet, H., Legendre, R., Pawlotsky, J.M., Sampaolesi, M. et al. (2021) Tissue damage induces a conserved stress response that initiates quiescent muscle stem cell activation. Cell Stem Cell, 28, 1125–1135 e1127.

50. Lowndes, M., Rakshit, S., Shafraz, O., Borghi, N., Harmon, R.M., Green, K.J., Sivasankar, S. and Nelson, W.J. (2014) Different roles of cadherins in the assembly and structural integrity of the desmosome complex. J Cell Sci, 127, 2339–2350.

51. Boivin, F.J. and Schmidt-Ott, K.M. (2017) Transcriptional mechanisms coordinating tight junction assembly during epithelial differentiation. Ann N Y Acad Sci, 1397, 80–99.

52. Schmelzer, C.E.H. and Duca, L. (2021) Elastic fibers: formation, function, and fate during aging and disease. FEBS J.

53. Brocker, C.N., Vasiliou, V. and Nebert, D.W. (2009) Evolutionary divergence and functions of the ADAM and ADAMTS gene families. Hum Genomics, 4, 43–55.

